# Continuous-flow synthesis and functionalization of gold nanoparticles for diagnostics and therapy

**DOI:** 10.64898/2026.11.20.768139

**Authors:** Demian Pardo, Francisco Tato, Carmen Escalona-Noguero, María López-Valls, Ciro Rodríguez-Díaz, Eduardo García-Garrido, Paula Milán-Rois, Begoña Sot, Álvaro Somoza

## Abstract

Gold nanoparticles (AuNPs) are highly versatile platforms in nanomedicine, but their clinical translation is hindered by poor reproducibility and scalability during synthesis and surface functionalization, which are still predominantly performed via manual batch protocols. To overcome these manufacturing bottlenecks, this work introduces a modular, continuous-flow millifluidic platform that integrates the automated synthesis of citrate-stabilized AuNPs and their subsequent multi-step surface modification within a single, streamlined pipeline. AuNPs of approximately 13 and 22 nm were reliably produced by adjusting the reagent stoichiometry. The flow-synthesized nanoconjugates showed hydrodynamic diameters comparable to those of the batch formulations, and comparable zeta potentials except for the PEG-containing ones. The platform’s chemical flexibility was validated through two distinct surface modifications: an in-line covalent ligand exchange mediated by bidentate dithiolane anchors, and a salt-free, ultra-fast low-pH adsorption method. To benchmark diagnostic performance against established batch protocols, flow-modified AuNPs were evaluated in previously optimized SARS-CoV-2 RNA sensing systems, a cholesterol-driven molecular beacon assay and the CRISPR-Cas13a-mediated CASCADE platform, reproducing the response time, specificity, and colorimetric readout of their batch-produced counterparts without requiring tedious salt-aging steps. The same hairpin design was also adapted to a lateral flow format. For therapeutic purposes, two distinct classes of bioactive agents were evaluated: a small-molecule chemotherapeutic system co-loaded with SN38 and the nucleolin-targeting AS1411 aptamer, which exhibited potent anti-proliferative activity against triple-negative breast cancer cells (MDA-MB-231), and a nucleic acid-based platform functionalized with the antisense oligonucleotide ASO34a via the low-pH module, which triggered significant post-transcriptional gene silencing (achieving at least 30% luciferase knockdown at 24 hours in both breast and pancreatic PANC-1 cancer cell lines). This automated millifluidic platform demonstrates that complex, multi-component nanomaterial modifications can be successfully executed under continuous-flow configurations, reproducing the performance of established batch-synthesized diagnostic assays and yielding active therapeutic nanoconjugates. By coupling precise core synthesis with sophisticated surface engineering, this continuous manufacturing paradigm establishes a scalable and reproducible pathway to narrow the gap between laboratory-scale discovery and clinical manufacturing.

## Introduction

Over the last few decades, the field of gold nanotechnology has undergone remarkable growth, driving a multitude of AuNP-based innovations from academic discovery toward clinical applications in diagnostics, biosensing, targeted therapy, and gene delivery^1,2^. These technologies have successfully transitioned into the commercial market as medical devices, particularly as colorimetric lateral flow assays (LFAs) for detecting critical cardiac biomarkers (such as Troponin I) and widespread infectious diseases (including SARS-CoV-2 and Malaria). Simultaneously, the clinical landscape of AuNPs has progressed from early safety assessments to sophisticated efficacy evaluations in human trials, with over twenty investigated formulations^3^. This clinical pipeline showcases a highly diverse range of molecularly modified AuNPs, including landmark targeted drug delivery systems carrying recombinant proteins like CYT-6091 (Aurimune) for the treatment of advanced solid tumors, spherical nucleic acid architectures functionalized with siRNA oligonucleotides like NU-0129 designed to target glioblastoma multiforme, multi-valent peptide-conjugated vaccines such as PepGNP-Dengue for infectious disease immunization, and even naked, clean-surface nanocrystals acting as metabolic catalysts for neurodegenerative disorders like amyotrophic lateral sclerosis (ALS) and multiple sclerosis (MS) (CNM-Au8)^3^.

Nevertheless, the translation of these highly specialized products from the laboratory bench to industrial and clinical settings remains heavily burdened by several critical challenges. Beyond the intrinsic biological and physicochemical complexities, such as precise structural characterization, long-term colloidal stability, controlled drug release, pharmacokinetics, and systemic toxicology, the core obstacle lies in the fact that the physicochemical properties of gold nanoparticles are strictly size- and shape-dependent^4^. Because these attributes are highly sensitive to even minor fluctuations in reaction parameters, any morphological deviation can profoundly alter their performance in nanomedicine applications. Furthermore, the interaction of these nanomaterials with biological structures can completely change, and potentially trigger unexpected systemic toxicity, depending on subtle variations in their dimensions and surface architecture^5^.

Conventionally, the synthesis of these nanomaterials has been performed in flask-type reactors due to its simplicity and straight-forwardness, which perfectly fits academic research purposes. However, the scaling up of batch-type reactors has been highly problematic; the inherent decrease in mixing efficiency and heat transfer within the reaction volume hinders proper control of the process^6^. In addition, the synthetic conditions for certain gold nanoparticles require harsh environments, such as elevated temperatures, high pressures, or extreme pH levels, raising significant safety concerns for both the operator and the environment^7^. Due to this high sensitivity and the limitations of traditional synthesis, the nanomedicine field faces severe industrial bottlenecks regarding batch-to-batch reproducibility and large-scale manufacturing^8–10^. Consequently, satisfying the rigorous multi-jurisdictional regulatory frameworks imposed by agencies such as the US Food and Drug Administration (FDA) and the European Medicines Agency (EMA) introduces a formidable hurdle. These authorities demand strict process validation, absolute structural homogeneity, and full compliance with evolving continuous manufacturing guidelines before granting clinical certification^11–13^.

In this sense, flow chemistry has arisen as a unique tool that offers exceptional design plasticity, stable operating production, and the ability to monitor processes safely. Continuous flow synthesis of nanometric materials allows for the spatial separation between the nucleation and growth phases within the same platform. In addition, flow reactors provide improved mixing rates alongside more efficient heat and mass transfer due to their large surface area-to-volume ratios compared to conventional batch-type reactors^6^. Thus, flow reactors represent highly promising tools for fine-tuning the size and morphology of nanomaterials in a reproducible manner. Consequently, the synthesis of AuNPs using continuous-flow reactors has emerged as a major trend in nanomanufacturing, driving intense research into its viability and scale-up potential within the framework of classical synthetic methods^14–19^. Among these, AuNPs synthesized via the Turkevich method represent a highly attractive option for nanomedical purposes due to the low intrinsic toxicity of sodium citrate. Building upon this method, recent research has focused on optimizing different aspects of the reaction. For instance, several studies have improved reagent mixing capacities within fluidic systems by incorporating innovative mixer designs^20–22^. Other publications have demonstrated that continuous flow can overcome the traditional size limitations of the Turkevich method, yielding AuNPs smaller than 10 nm with remarkably low heterogeneity^23,24^. Furthermore, the reproducibility and scaling of the process have been extensively analyzed. Panariello et al. investigated methods to enhance synthetic reproducibility by evaluating the effect of passivating the gold precursor stock solution prior to its use^25^. Their findings showed that utilizing a pre-passivated precursor provided superior control over the initial reaction stage, albeit at the expense of a slight increase in reaction time. Expanding on process robustness, Damilos et al. developed a setup that allowed the stable synthesis of 10–20 nm AuNPs over a 2-hour continuous run, achieving yields between 70% and 100% depending on the flow rate^26^. Beyond core synthesis, the in-line modification of AuNPs inside flow reactors has also been successfully accomplished. For example, Gomez et al. performed the single-step functionalization of hollow gold nanoparticles with thiolated PEG to enhance nanomaterial stability and extend its biological half-life^27^. Similarly, Uson et al. leveraged a thiolated PEG ligand exchange strategy to substitute the cytotoxic surfactant CTAB with a biocompatible polymer on gold nanorods (GNRs)^28^.

In addition to polyethylene glycol (PEG) shielding, the true translational power of AuNPs lies in their capacity to be co-conjugated with a vast and diverse array of biological, therapeutic, and targeting molecules. For instance, when densely functionalized with DNA or RNA oligonucleotides, AuNPs form the so-called “Spherical Nucleic Acids” (SNAs), exhibiting exceptional stability, high affinity for complementary sequences, and an inherent capacity for efficient cellular internalization^29,30^. However, the scope of AuNP hybridization extends far beyond polymer stabilization and nucleic acids alone; the precise integration of small-molecule chemotherapeutics alongside highly specific targeting ligands, such as aptamers, allows for the engineering of sophisticated, multipurpose nanomedicines capable of simultaneous targeting, therapy, and diagnosis^31–34^. Herein, we assess the viability of continuous-flow reactors for both the synthesis and multi-step functionalization of AuNPs with nucleic acids and other diverse therapeutic molecules tailored for the nanomedicine field. In this work, AuNPs are sequentially modified with nucleic acids, PEG, and the small-molecule drug SN38 using a robust, in-line covalent strategy within a flow platform. Subsequently, the biomedical functionality of these flow-modified AuNPs is validated through their application as optical biosensors for SARS-CoV-2 diagnosis, as well as targeted drug delivery agents for cancer treatment and gene silencing. Ultimately, this study demonstrates that automated flow platforms enable the precise, in-line modification of AuNPs with high-value biomolecules, highlighting their potential translation into commercial market realities. To the best of our knowledge, this represents the first continuous-flow platform that integrates, within a single in-line process, the automated synthesis and multi-step covalent functionalization of AuNPs with biomolecules as diverse as nucleic acids, PEG, a small-molecule drug, and an aptamer, effectively bridging advanced diagnostic and therapeutic applications into a feasible nanomanufacturing platform.

## Materials and Methods

### Materials and reagents

Trisodium citrate dihydrate (C6H5Na3O7)•2H2O, citric acid monohydrate (C6H8O7)•H2O, and sodium chloride (NaCl) were purchased in powder form from Sigma-Aldrich. Hydrogen tetrachloroaurate(III) trihydrate (HAuCl4)•3H2O was obtained from Alfa Aesar. HEPES (4-(2-hydroxyethyl)-1-piperazineethanesulfonic acid) and Gibco™ Opti-MEM™ I Reduced Serum Medium were purchased from Fisher Scientific. Pure silicone liquid was sourced from Scharlau. Ultra-pure water (Milli-Q) was obtained using a Milli-Q Advantage A10 purification system. Sample pads, absorbent pads (C083), and glass fiber conjugate pads (G041) were purchased from Sigma-Aldrich, while nitrocellulose membranes (FF80HP) were obtained from Cytiva. DMEM High Glucose and RPMI 1640 cell culture media, the alamarBlue cell viability assay kit, and Lipofectamine 2000 and 3000 transfection reagents were purchased from Thermo Fisher Scientific (Invitrogen). Corning 96-well clear flat-bottom plates, black 96-well plates (for fluorescence assays), and white 96-well microplates (for luminescence measurements) were obtained from VWR. The Dual-Luciferase Reporter Assay System was purchased from Promega. All other chemical reagents were of analytical grade and used as received without further purification.

### Nucleic Acid Sequences and origin

Oligonucleotides were either purchased from Integrated DNA Technologies (IDT) or synthesized using H8/H6 DNA/RNA automated synthesizers (K&A Labs GmbH). Following solid-phase synthesis, the solid support was transferred to a screw-cap glass vial and cleaved/deprotected by incubating at room temperature for 18 h with 2 mL of an aqueous ammonia solution (32%) for DNA, or an ammonia/ethanol mixture (3:1, v/v) for RNA. Subsequently, the crude samples were dried and purified using standard DNA/RNA purification columns (LCG Biosearch Micropure purification columns). After purification, the samples were dried once more and resuspended in 200 µL of RNAse-free water. Nucleic acid concentrations were determined via UV-Vis spectrophotometry by measuring the absorbance at 260 nm using a Cary 5000 UV-Vis-NIR Spectrophotometer (Agilent Technologies). Molar extinction coefficients were calculated utilizing the online OligoAnalyzer™ software tool from IDT. The names, specific sequences, and experimental targets of all nucleic acids utilized for both batch and continuous-flow functionalization of AuNPs are detailed in Table 1.

**Table 1.**
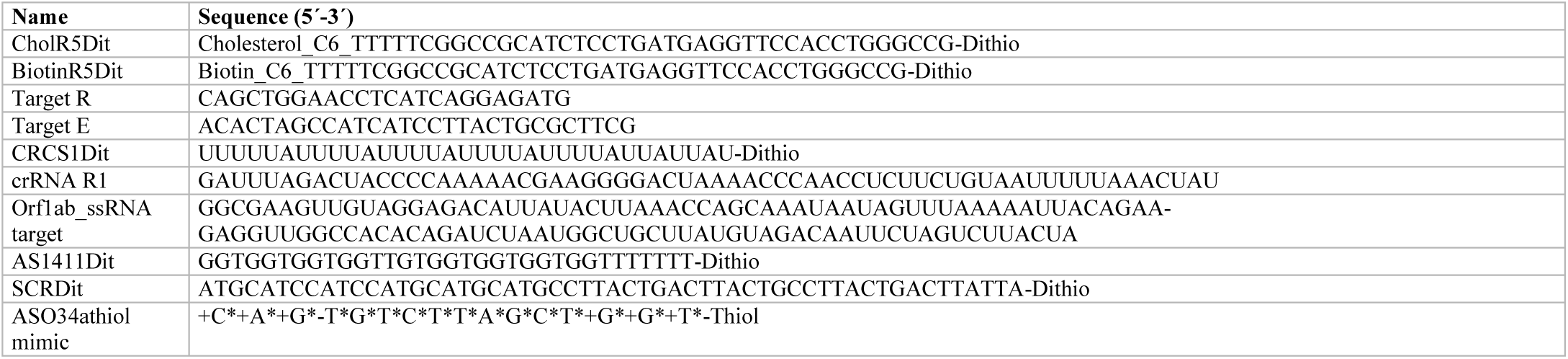
Nucleic acids used during the functionalization of AuNPs in the flow reactor, batch conditions, and targets. += Locked nucleic acid; *= Phosphorothioate

### Synthesis of dithiolane-modified PEG (PEG-Dithiolane)

A 2 kDa poly(ethylene glycol) (PEG 2K) functionalized with a dithiolane linker was prepared. Briefly, this conjugate was synthesized via the covalent coupling of lipoic acid and PEG 2K in anhydrous tetrahydrofuran (THF) using N-hydroxysuccinimide (NHS) and N,N-dicyclohexylcarbodiimide (DCC) as standard coupling reagents, following previously reported protocols^35^.

### Synthesis of dithiolane-modified SN38 (SN38-Dithiolane)

A modified hydroxycamptothecin derivative bearing a dithiolane linker was prepared. The conjugate was synthesized by coupling lipoic acid with the chemotherapeutic drug SN38 in dimethyl sulfoxide (DMSO) using NHS and DCC as standard coupling reagents^36^.

### Construction and purification of the pmirGLO-miRNA34a plasmid

The pmirGLO Dual-Luciferase miRNA Target Expression Vector was linearized and subsequently ligated with miRNA34a oligonucleotides according to the manufacturer’s instructions. The resulting ligated pmirGLO-miRNA34a vector was transformed into chemically competent *E. coli* DH5α cells. Positive clones were selected on agar plates supplemented with ampicillin. Plasmid extraction and purification were performed using the NZY Miniprep kit, and the final concentration of the purified miRNA34a plasmid was quantified using a NanoDrop spectrophotometer.

### CRISPR-Cas 13 complex

#### Protein expression and purification of LwaCas13a

LwaCas13a was expressed and purified by adapting a previously described protocol^37^. Briefly, the bacterial expression vector p2CT-His-MBP-Lwa_Cas13a_WT (Addgene) was transformed into competent *Escherichia coli* BL21 cells. A starter culture (25 mL) was grown overnight in Luria-Bertani (LB) broth and subsequently used to inoculate 3 L of LB medium supplemented with ampicillin. Cells were cultured at 37°C with shaking at 200 rpm until reaching an optical density (OD600) of 0.3. The temperature was then shifted to 16°C, and protein expression was induced with 0.5 mM isopropyl β-D-1-thiogalactopyranoside (IPTG) once the (OD600) reached 0.5–0.6. After overnight incubation, biomass was harvested by centrifugation, resuspended in lysis buffer (50 mM Tris-HCl pH 7.5, 500 mM NaCl, 5% glycerol, 5 mM β-mercaptoethanol, 0.5 mM PMSF, lysozyme, and EDTA-free protease inhibitor), and disrupted via sonication. The lysate was cleared by centrifugation at 10,000 rpm for 1 h at 4°C. Soluble His6-MBP-TEV-Cas13a was captured using a HisTrap HP column (GE Healthcare). Target protein fractions were pooled, concentrated, and incubated overnight at 4°C with TEV protease during dialysis against a buffer containing 20 mM Tris-HCl pH 7.0, 500 mM NaCl, 5% glycerol, and 1 mM DTT. The cleaved protein was further purified by size-exclusion chromatography on a Superdex 200 column (GE Healthcare). Finally, DTT was removed using a Nap-10 desalting column (GE Healthcare) equilibrated with 20 mM Tris-HCl pH 7.0, 500 mM NaCl, and 5% glycerol. The purified LwaCas13a was concentrated, aliquoted, and stored at -80°C. All purification steps were carried out at 4°C.

#### crRNA design

The crRNA R1 targeting LwaCas13a was designed to target RNA regions with minimal secondary structure, as previously described by Zhang et al.^38^. Secondary structure predictions for both the crRNAs and their corresponding targets were performed using UNAFold (IDT) and RNAfold (Vienna RNA WebServices) software. Modified or synthetic crRNAs were synthesized by Integrated DNA Technologies (IDT).

#### Target RNA preparation

The SARS-CoV-2 Orf1ab RNA target was generated via RT-PCR from total RNA extracted from infected cells (kindly provided by Dr. Isabel Sola and Dr. Sonia Zúñiga, CNB-CSIC), using the SuperScript™ III OneStep RT-PCR System with Platinum™ Taq DNA Polymerase (Thermo Fisher Scientific). Primers containing the T7 promoter sequence at the 5’ end were synthesized according to Zhang et al.^38^. *In vitro* transcription was performed using the mMESSAGE mMACHINE™ or MEGAscript™ T7 Transcription Kit (Thermo Fisher Scientific) for 16 h at 37°C. Following transcription, template DNA was degraded with TurboDNase for 25 min at 37°C. The synthesized RNA was purified via phenol-chloroform extraction: samples were diluted with 115 µL of RNAse-free water and 15 µL of Ammonium Acetate stop solution, followed by sequential extractions with phenol and chloroform. RNA was precipitated with ice-cold absolute ethanol and incubated at −20°C for 2 h. Precipitated RNA was recovered by centrifugation at maximum speed for 20 min at 4°C. The supernatant was removed, the pellet was air-dried to eliminate residual ethanol, and the final RNA target was resuspended in RNAse-free water.

#### Lateral flow assay strip fabrication

The lateral flow strip sensors were composed of a sample pad, a conjugation pad, a test line, and an absorbent pad. For strip pre-treatment, the sample pads were incubated overnight at 37°C in 1X PBS containing 0.1% PEG 3K. Concurrently, the conjugate pads were incubated in a blocking solution composed of 4% BSA, 0.5% Tween 20, 5% trehalose, and 0.05% sodium azide.

Immobilization of streptavidin onto the test line area was performed using a two-step crosslinking procedure adapted from a previously described method^39^. First, 1,4-diaminobutane (1 mg/mL) in RNAse-free water was deposited onto the target area for 1 h at room temperature, followed by two 1 h washing steps with RNAsefree water. Next, glutaraldehyde (0.5%) in RNAse-free water was applied for 15 min at room temperature, followed by two 30 min washes with RNAse-free water. Subsequently, streptavidin (10 mg/mL) dissolved in a buffer containing 1X PBS, 0.5% Tween 20, and 30% NaCl (0.5 M) was added and incubated for 6 h at room temperature. Finally, the functionalized strips were washed twice with a solution of 10 mM Tris-HCl (pH 8) and 0.1% Tween 20.

#### Cell lines

The triple-negative breast cancer cell line MDA-MB-231 was kindly provided by Dr. Ana Pizarro’s group (IMDEA Nanociencia, Spain), originally purchased from the American Type Culture Collection (ATCC). The pancreatic cancer cell line PANC-1 was purchased directly from the ECACC.

## Methods

### Experimental set- up

#### First stage: gold nanoparticle synthesis

The initial section of the flow reactor designed for the synthesis of AuNPs was identical across all the experiments. In detail, HPLC pumps (ProStar 215 Solvent Delivery Module, Varian) were employed to inject the reagents at a flow rate of 500 μL/min each. Two PTFE heating coils (1.6 mm OD, 1 mm ID, 0.1 meter; Vici) were utilized to preheat the reagent streams prior to mixing in a PEEK T-junction mixer. The resulting mixture was directed into a main PTFE flow reactor (3.2 mm OD, 1.6 mm ID, 10 m; Polyfluor) providing a residence time of 20 min. The entire reactor assembly was immersed and heated in a silicone oil bath maintained at 100°C. To ensure stable flow conditions, a backpressure regulator (40 PSI; Idex Health & Science) was positioned at the reactor outlet. Finally, a cooling coil (3.2 mm OD, 1.6 mm ID, 10 m; Vici) was incorporated to allow the synthesized AuNPs to reach room temperature. A schematic representation of this first stage of the system is illustrated in **Figure 1**.

**Figure 1.**
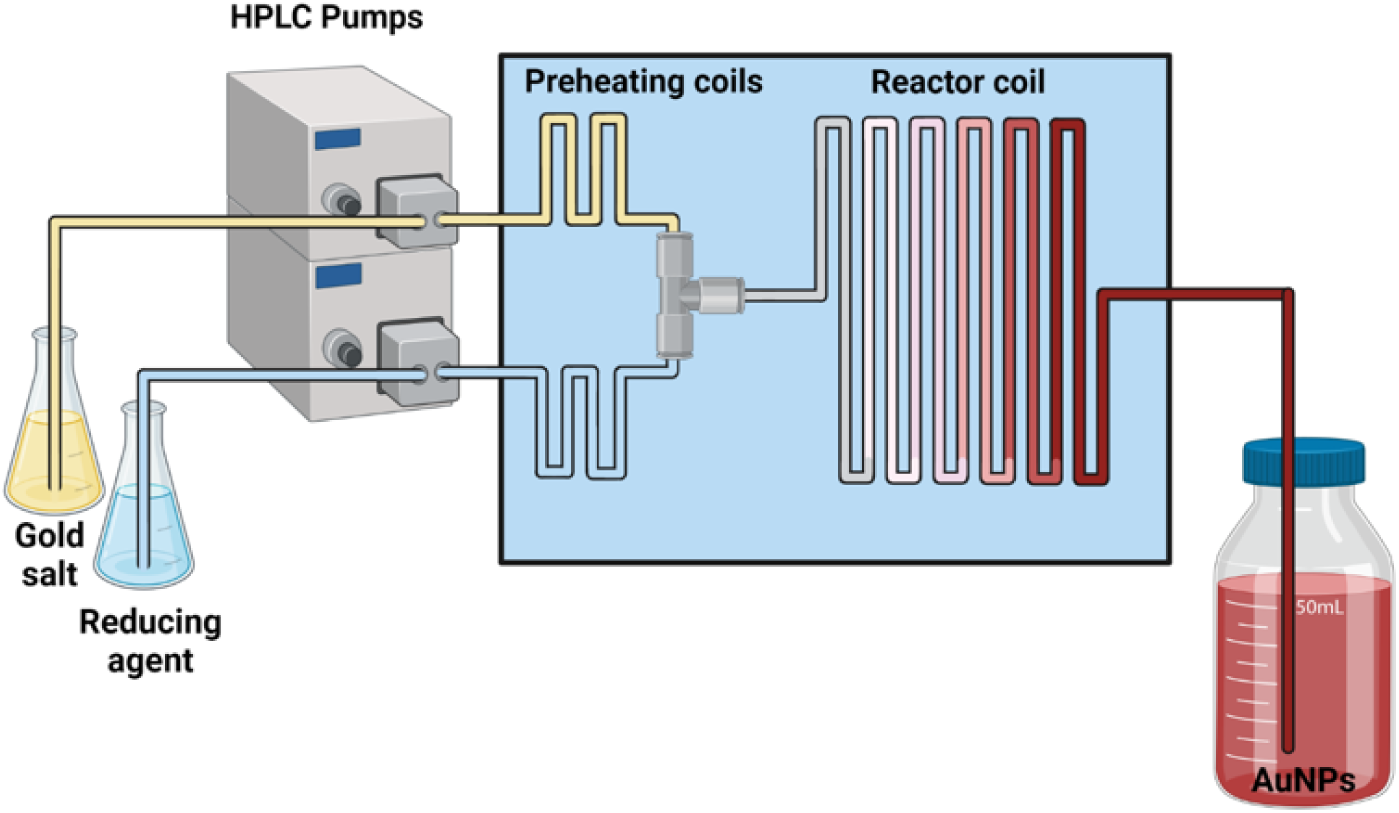
Schematic representation of the continuous-flow reactor setup designed for the synthesis of gold nanoparticles.

The specific amounts of gold salt and trisodium citrate, each dissolved in 55 mL of ultra-pure water and delivered by the HPLC pumps are summarized in **Table 2**.

**Table 2.** Amounts of gold salt and trisodium citrate (each dissolved in 55 mL of ultra-pure water) used across the different functionalization experiments.

| **Experiment** | **Gold salt (mmol)** | **Trisodium citrate (mmol)** | **Size (nm)** |
|---|---|---|---|
| Detection assay of SARS-CoV-2 with AuNPs functionalized with a cholesterol molecular beacon | 0.025 | 0.275 | 22 |
| Detection assay of SARS-CoV-2 with AuNPs functionalized with a biotin molecular beacon | 0.025 | 0.275 | 22 |
| Detection assay of SARS-CoV-2 with AuNPs functionalized with RNA by CRISPR-Cas13a | 0.1 | 0.7 | 13 |
| Cell viability assay with AuNPs functionalized with nucleic acids and SN38 | 0.1 | 0.7 | 13 |
| Gene silencing assay with AuNPs functionalized with ASO34a mimic | 0.1 | 0.7 | 13 |

### The second stage: the functionalization modules

The second section of the flow reactor was designed modularly for the surface functionalization of the synthesized AuNPs. This stage incorporated between one and four subsequent reaction modules, depending on the number of molecules required for each specific functionalization experiment, each equipped with its respective syringe pump. Specifically, the room-temperature AuNPs solution leaving the cooling coil was directed into the first custom-built PEEK dynamic mixer, where the first functionalization agent was injected via a syringe pump (SyringeONE Programmable Syringe Pump, model NE-1010). The mixture then entered a PTFE flow reactor (3.2 mm OD, 1.6 mm ID, 10 m; Polyfluor) providing a residence time of approximately 20 min.

Depending on the number of sequential addition steps required, up to three additional PEEK mixers were integrated in series. Each mixer featured an inlet connected to a dedicated syringe pump (SyringeONE Programmable Syringe Pump, model NE-1010). Downstream of each addition point, a PTFE flow reactor (3.2 mm OD, 1.6 mm ID, 10 m; Vici) was installed to afford a residence time of around 20 min. As an exception, for the surface modification of AuNPs with thiol-terminated molecules, three consecutive PTFE flow reactors (3.2 mm OD, 1.6 mm ID; Vici) of varying lengths were utilized along with their corresponding mixers to maintain a constant nominal residence time of 2 min per step. Specifically, the length of the first, second, and third reactors was adjusted to 1.12 m, 1.34 m, and 2.34 m, respectively, to precisely compensate for the increasing total flow rate. A schematic diagram and a photograph of the complete flow reactor configured with four addition steps are presented in **Figure 2** and **Figure 3**, respectively.

**Figure 2.**
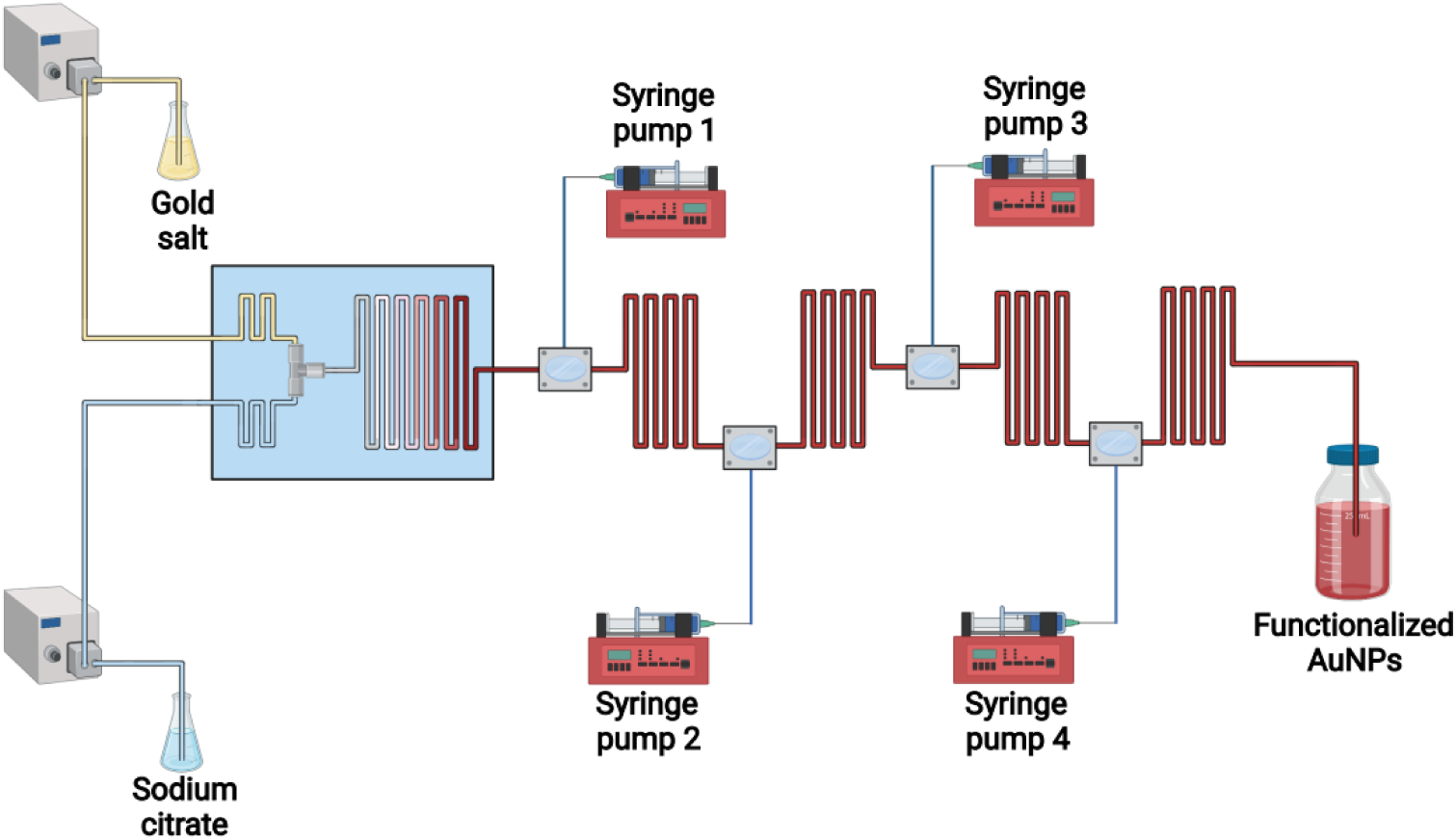
Complete schematic diagram of the modular continuous-flow reactor configured with four sequential functionalization steps.

**Figure 3.**
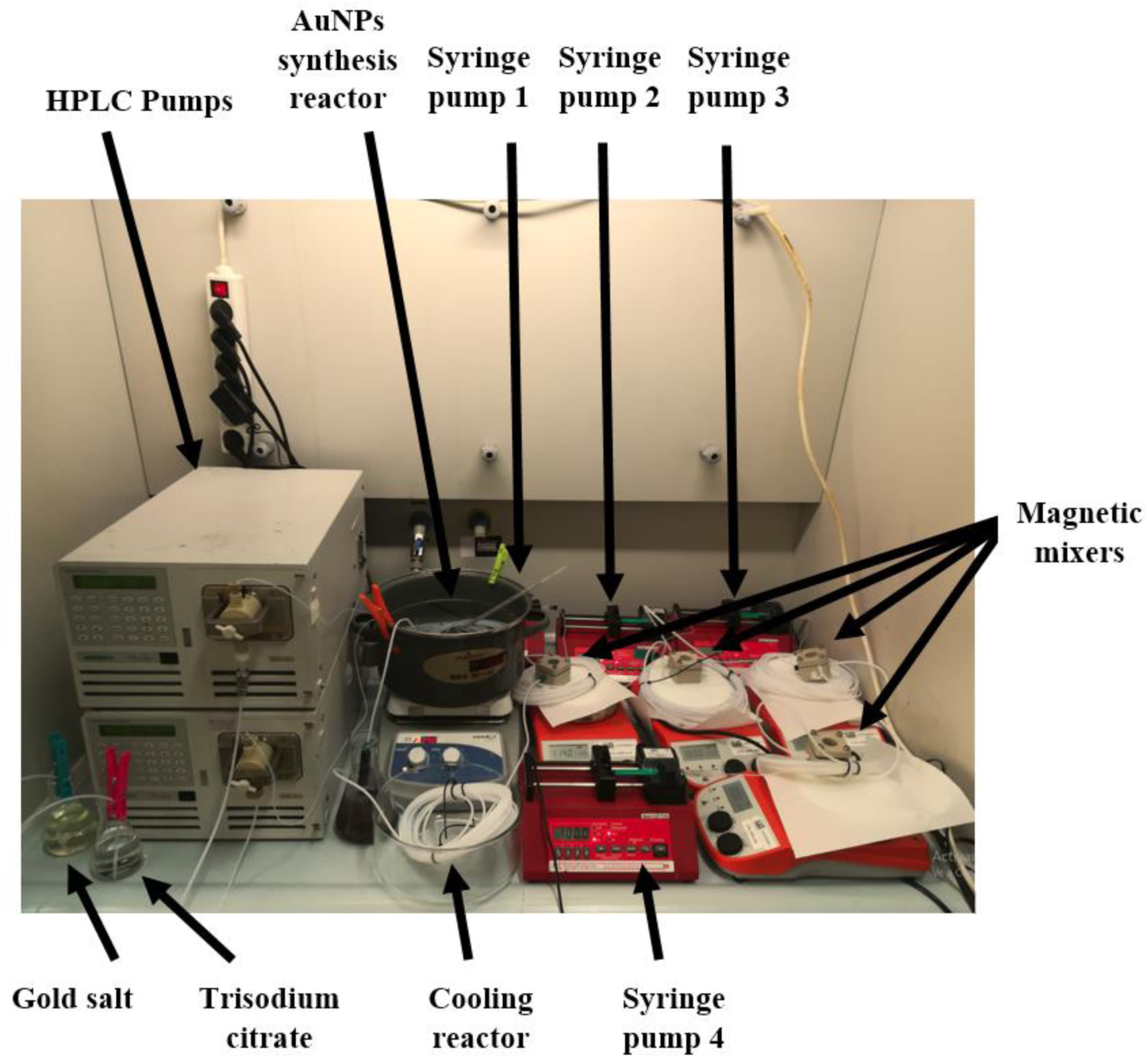
Photographic view of the complete continuous-flow reactor setup configured with four sequential functionalization steps.

Importantly, after each functionalization experiment, the PTFE tubing was cleaned with aqua regia. Prior to each functionalization experiment, the full setup was cleaned by flushing an aqueous solution of EDTA and NaOH, followed by a thorough rinse with RNAse-free water to eliminate trace metal contaminants and nucleases.

The precise addition sequence, reagent concentrations, and operational parameters for each individual experiment are summarized in **Supplementary Table 1**.

### Post-synthetic treatments

Once functionalized, all AuNPs were washed three times by centrifugation at 10,000 rpm for 20 min at 4 °C. The 13 nm AuNPs were resuspended to their original volume in RNAse-free water. The 22 nm AuNPs were concentrated up to 6.37 nM for the SARS-CoV-2 molecular beacon detection assay, whereas the 13 nm RNA-functionalized AuNPs were concentrated to 23.4 nM for the CRISPR-Cas13a SARS-CoV-2 detection assay.

### Characterization of gold nanoparticles

The morphology and dimensions of the synthesized AuNPs were evaluated using Transmission Electron Microscopy (TEM) and Atomic Force Microscopy (AFM). TEM imaging was conducted at the CBMSO-CSIC facilities using a JEM1400 Flash microscope (Jeol, Japan) operated at 120 keV. For sample preparation, a drop of the AuNP suspension was deposited onto a glow-discharged, Formvar-coated, carbon-shadowed HEX 400-mesh copper grid (Agar Scientific) for 2 min, followed by the removal of excess liquid. Size distributions and nanoparticle shapes were quantified via automated image analysis of random fields using ImageJ software, measuring a minimum of 250 AuNPs per experimental condition. Complementarily, the concentration and optical size of the nanoparticles were assessed from UV–vis absorption spectra acquired with a Cary 5000 Spectrophotometer (Agilent Technologies), following a previously reported methodology^40^. Specifically, the core diameter was estimated from TEM micrographs and the ratio between the localized surface plasmon resonance (LSPR) peak absorbance and the absorbance at 450 nm. This calculated diameter was subsequently used to determine the nanoparticle concentration by applying the corresponding extinction coefficient at 450 nm.

### Detection assay of SARS-CoV-2 with AuNPs functionalized with a cholesterol molecular beacon

To evaluate the diagnostic performance of the AuNPs functionalized with CholR5Dit within the flow reactor, the nanoconjugates were incubated with a specific synthetic target RNA sequence (R) corresponding to SARS-CoV-2. The reaction kinetics were monitored visually by capturing digital images at 0, 1, and 2 h. Concurrently, aliquots were transferred to a 96-well UV-vis plate at the same time points to record the absorbance spectra from 400 to 650 nm (3 nm step size) using a Synergy H4 Plate Reader.

### Detection assay of SARS-CoV-2 with AuNPs functionalized with the biotin molecular beacon

To assess the diagnostic performance of the flow-synthesized AuNPs functionalized with BiotinR5Dit for SARS-CoV-2 detection, lateral flow assays were performed. The functionalized AuNPs were incubated with varying concentrations of target R (1000, 500, 250, 100, and 25 nM) dissolved in a running buffer composed of 4% BSA, 0.5% Tween 20, 5% trehalose, and 0.05% sodium azide. The mixtures were allowed to react for 10 min at room temperature to ensure proper binding before being loaded onto the sample pad of the prepared test strips. Following completion of the lateral flow run, the strips were rinsed with RNAse-free water to eliminate non-specific background signals. Digital photographs of the strips were captured using a standard camera. Additionally, high-resolution optical densitometry images of the Ponceau S-treated strips were acquired using a ChemiDoc Imaging System and subsequently quantified using ImageJ software.

### Detection assay of SARS-CoV-2 with AuNPs functionalized with RNA by CRISPR-Cas13a

To evaluate the diagnostic capability of the AuNPs functionalized with CRCS1Dit within the flow reactor, 50 μL of the functionalized nanoconjugates (23.4 nM) were mixed with 50 μL of Cas13a ribonucleoprotein (RNP, 50 nM) diluted in cleavage buffer containing 60 mM Tris-HCl, 60 mM NaCl, and 6 mM MgCl₂. Subsequently, the target RNA was added at a final concentration of 25 nM, and the mixtures were incubated at 37°C. The reaction kinetics were visually monitored by capturing digital images at 0, 15, 30, and 60 min. Concurrently, aliquots were transferred to a 96-well UV-plate at the same time points to record the absorbance spectra from 400 to 650 nm (3 nm step size) using a Synergy H4 Plate Reader.

### Cell viability assay with AuNPs functionalized with nucleic acids and SN38

To evaluate whether nucleic acids enhance the delivery of SN38, various functionalized AuNPs were synthesized in the flow reactor, including AuNPs modified with SN38Dit + PEGDit, SN38Dit + AS1411Dit, SN38Dit + SCRDit, as well as the controls AuNPs + PEGDit and AuNPs + SCRDit. The therapeutic efficacy of these flow-synthesized nanoconjugates was assessed via a cell viability assay using the triple-negative breast cancer cell line MDA-MB-231. Cells were cultured in DMEM High Glucose medium supplemented with 10% fetal bovine serum (FBS), 1% penicillin-streptomycin, and 1% L-glutamine at 37°C in a humidified atmosphere containing 5% CO₂ using a Binder CB210 incubator. All cell culture procedures were conducted inside a Bio II Advance laminar flow hood (Telstar, Terrassa, Spain). For the cytotoxicity assays, MDA-MB-231 cells were seeded in 96-well plates at a density of 10,000 cells/well and grown until achieving 60–70% confluence. Prior to treatment, the respective AuNP formulations were concentrated to 150 μL and diluted in supplemented DMEM High Glucose medium to a final volume of 600 μL. Cells were then exposed to each experimental condition in hexaplicate (n = 6). Metabolic activity was quantified using resazurin sodium salt (Alamar Blue assay) dissolved in PBS (1 mg/mL). Briefly, the culture medium was replaced with 100 μL of a 1% (v/v) resazurin solution in fresh medium, followed by incubation at 37°C for 3 h. For fluorometric analysis, 80 μL aliquots from each well were transferred to a black 96-well plate. Fluorescence intensity was measured at endpoint after 24, 48, and 72 h at 25°C using a Synergy H4 Plate Reader, with excitation and emission wavelengths set to 550 and 590 nm, respectively. A cell-free resazurin solution served as the negative control.

### Gene silencing assay with AuNPs functionalized with ASO34a mimic

To evaluate the gene silencing efficiency of flow-synthesized nanoconjugates, AuNPs functionalized with a miR-34a mimic (ASO34a-AuNPs) were prepared. Luciferase knockdown capability was assessed in the triple-negative breast cancer cell line MDA-MB-231 and the pancreatic cancer cell line PANC-1. Both cell lines were maintained at 37°C under a humidified 5% CO₂ atmosphere, using DMEM High Glucose medium for MDA-MB-231 and RPMI 1640 medium for PANC-1. Both media were supplemented with 10% fetal bovine serum (FBS), 1% penicillin-streptomycin, and 1% L-glutamine. All cell culture operations were performed inside a Bio II Advance laminar flow hood (Telstar, Terrassa, Spain). For both silencing and cytotoxicity experiments, cells were seeded in 96-well plates at a density of 10,000 cells/well and cultured until reaching 60–70% confluence. All treatments were performed in triplicate (n = 3). For the dual-luciferase reporter assay, cells were first transfected with the pmirGLO Dual-Luciferase miRNA34a plasmid vector for 4 h. Subsequently, the media were replaced, and cells were treated with ASO34a-AuNPs at final concentrations of 3, 6, and 9 nM. After 24 h of incubation, cell lysates (90 μL) were transferred to white 96-well luminescence plates. Bioluminescence endpoints for *Renilla* and firefly luciferases were sequentially recorded at 25°C using a Synergy H4 Plate Reader, with emission filters set at 470 nm and 560 nm, respectively. Concurrently, cell viability was evaluated using the resazurin sodium salt (Alamar Blue) assay. Cells were exposed to ASO34a-AuNPs (3, 6, and 9 nM) for 24 h. Afterwards, the culture medium was replaced with resazurin dissolved in PBS (1 mg/mL) at a final concentration of 1% (v/v) in fresh medium. Following a 3 h incubation at 37°C, metabolic activity was quantified by measuring fluorescence intensity at endpoint (λex = 550 nm, λem = 590 nm) at 25°C utilizing the Synergy H4 Plate Reader.

### Statistical Analysis

Statistical analyses were performed using GraphPad Prism software. Statistically significant differences between the mean values of the experimental conditions and the control group were evaluated via one-way analysis of variance (ANOVA), followed by Tukey’s post-hoc test among experimental groups. Differences were considered statistically significant at p < 0.05 (*), p < 0.01 (**), and p < 0.001 (***).

## Results

### Synthesis and characterization of flow-synthesized AuNPs

AuNPs can be precisely tuned to achieve specific dimensions and geometries tailored for diverse nanomedicine applications^41,42^. Here, the classical Turkevich method was adapted to the automated flow reactor to synthesize spherical AuNPs of approximately 13 and 22 nm in a straightforward and robust manner^42^. To optimize the dimensions, the influence of the stoichiometric ratio (R) between trisodium citrate and the gold precursor (_HAuCl4_) was systematically evaluated at R values of 3, 4, and 7.

Transmission electron microscopy (TEM) images revealed a progressive decrease in the average core diameter as the R ratio increased. Specifically, the mean sizes of the AuNPs synthesized at R = 3, R = 4, and R = 7 were 21 ± 3 nm, 16 ± 2 nm, and 13 ± 1 nm, respectively (**Figure 4**). Notably, the nanoconjugates obtained at R = 7 exhibited the highest size homogeneity, whereas lower ratios (R = 3 and R = 4) yielded more polydisperse populations. This trend was further supported by the calculated coefficients of variation (CV), which decreased from 16.6% to 12.7% and 10.0% as R shifted from 3 to 7, confirming that higher citrate concentrations favor the formation of more monodisperse and smaller metallic cores under continuous flow conditions.

**Figure 4.**
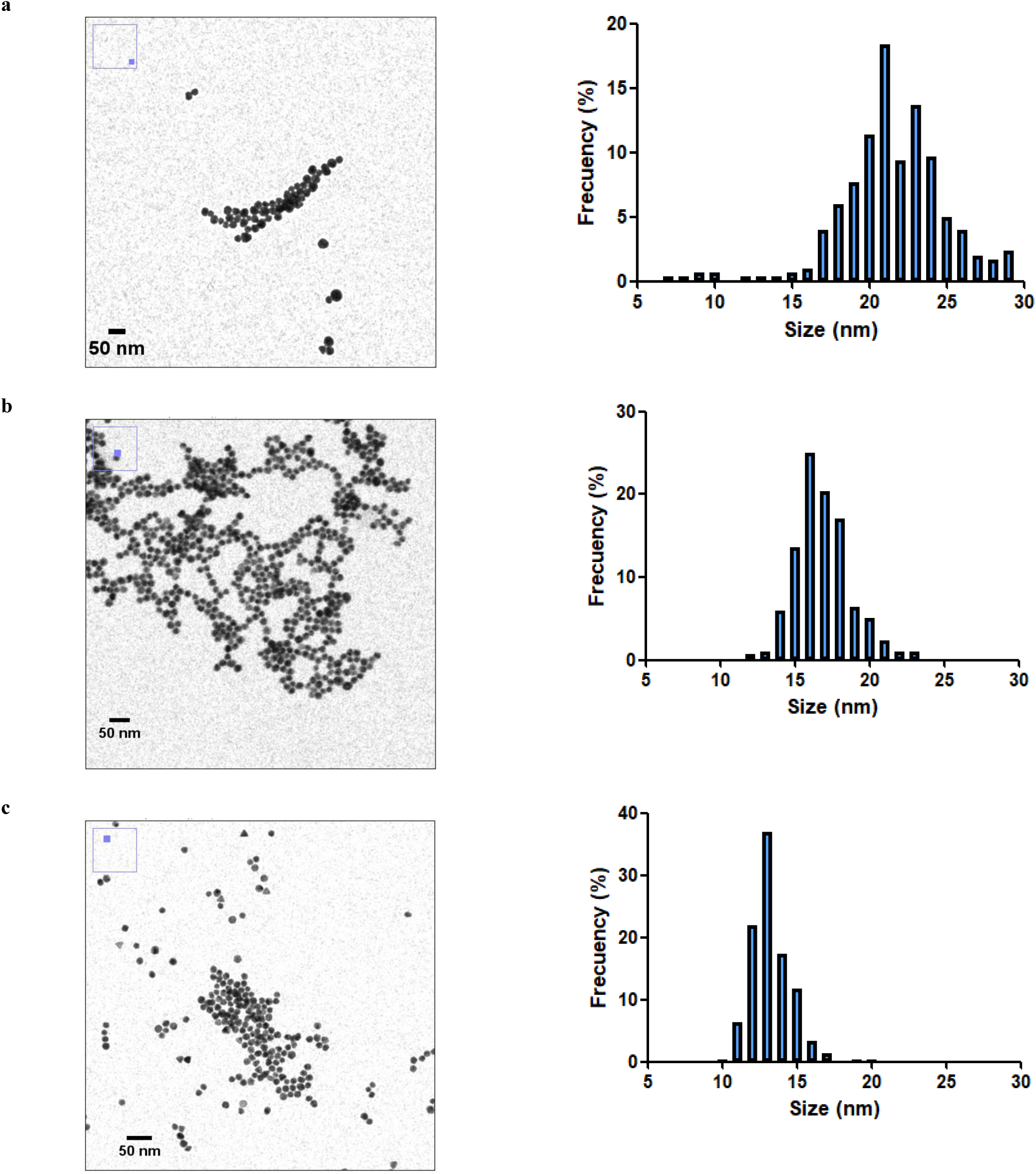
Transmission electron microscopy images and corresponding core size distributions of the synthesized AuNPs obtained at citrate-to-gold precursor molar ratios (R) of (a) R = 3, (b) R = 4, and (c) R = 7.

Although the AuNPs synthesized at R = 3 (ca. 21 nm) were within the target size range, the resulting pH of the solution was less than 4.0. This led to poor colloidal stability, which was directly attributed to the protonation of citrate molecules at low pH levels, reducing their electrostatic stabilization efficiency^43^. To overcome this limitation and achieve stable nanoparticles of larger dimensions, an alternative strategy within the Turkevich framework was implemented by increasing the sodium citrate concentration. At higher pH levels induced by an excess of citrate, the nucleation rate significantly decreases; this reduction results in a lower number of initial seed particles, which subsequently consume the remaining gold precursor to yield larger final core sizes^44^. Consequently, the molar ratio was increased to 11 (R = 11). Under these conditions, TEM images revealed the formation of stable AuNPs with an average core size of 22.2 ± 3.1 nm and a coefficient of variation (CV) of 13.7% (**Figure 5**).

**Figure 5.**
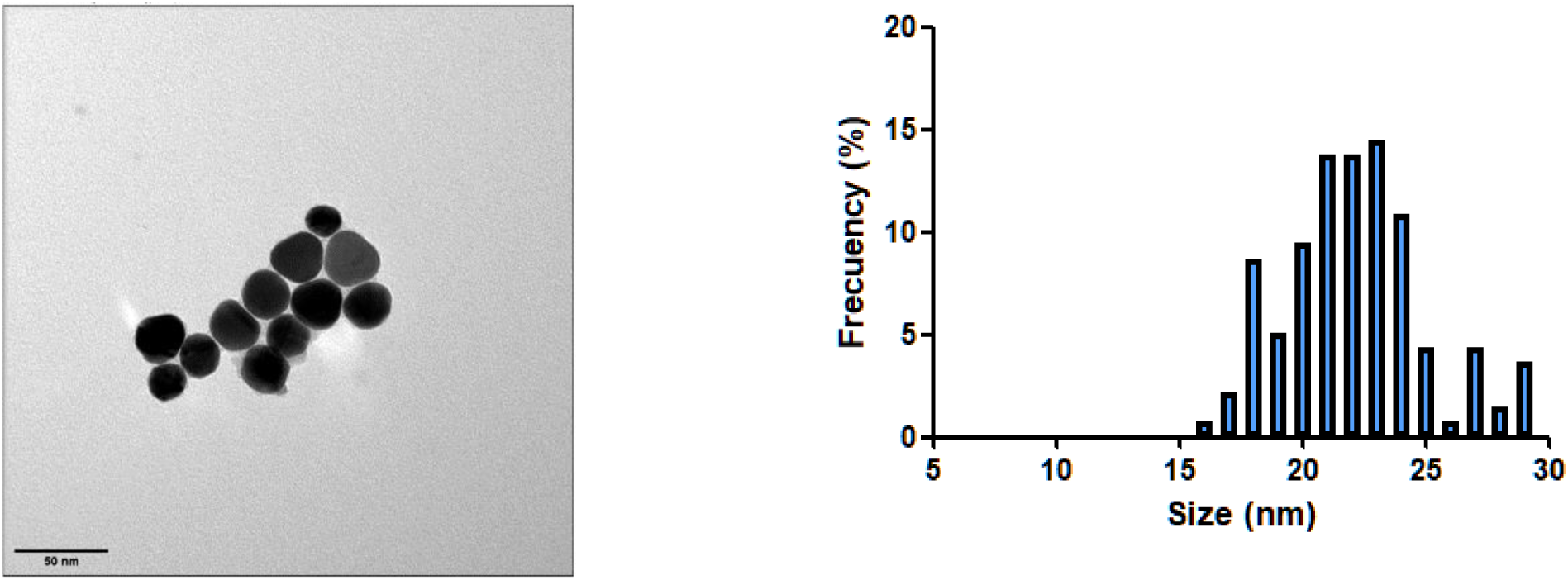
Transmission electron microscopy image and corresponding core size distribution of the synthesized AuNPs obtained at a citrate-to-gold precursor molar ratio of R = 11 (n = 250 nanoparticles measured per condition).

Thus, for the subsequent functionalization experiments, the initial continuous-flow synthesis of the gold nanoparticles was performed at R ratios of 7 and 11 to yield the 13 and 22 nm AuNP platforms, respectively.

### Biomedical validation of continuous-flow functionalized AuNPs

To demonstrate the translational potential and functional reliability of our continuous-flow platform, we evaluated the flow-synthesized nanoconjugates across both diagnostic and therapeutic applications. For the diagnostic systems, we built on two colorimetric SARS-CoV-2 RNA sensing strategies previously developed and optimized by our group using conventional batch protocols: a hydrophobic molecular beacon-based assay^45^ and the CRISPR-Cas13a-mediated CASCADE system^46^. By replicating the key proof-of-concept assays using flow-modified AuNPs, we aimed to confirm that automated continuous-flow functionalization preserves the physicochemical integrity and biosensing performance of the batch-produced benchmarks. The results obtained here are compared with those reported in those studies, so that the validation of the platform rests on a direct comparison with published examples. In addition, the same hairpin design was adapted to a lateral flow format by replacing the cholesterol with a biotin reporter. For the therapeutic applications, we selected two examples that use two types of therapeutic agents: a chemotherapeutic drug (SN38, co-functionalized with the targeting aptamer AS1411) and a nucleic acid-based therapy (the antisense oligonucleotide ASO34a for gene silencing).

### Diagnostic applications: SARS-CoV-2 RNA detection

#### Detection assay of SARS-CoV-2 with AuNPs functionalized with a cholesterol molecular beacon

Molecular beacons have emerged as a powerful strategy for the development of fluorescent-based AuNP sensors^47^. In a conventional design, the short-hairpin probe consists of approximately 25 nucleotides, where 10 to 15 bases form the loop region designed to be complementary to the target DNA/RNA sequence, while lacking self-complementarity. Conversely, five bases at each terminus are complementary to each other, forming a stem structure that maintains the fluorophore and the quencher in close spatial proximity. Upon encountering a sample containing the target sequence, hybridization occurs at the loop region. Because the number of nucleotides involved in this target hybridization outnumbers those stabilizing the stem, the structural equilibrium shifts, inducing the conformational opening of the hairpin. Consequently, the distance between the fluorophore and the quencher increases, leading to the restoration of the fluorescent signal.

Capitalizing on this mechanism, an alternative approach developed by Latorre et al. replaces the fluorescent reporter with a hydrophobic cholesterol moiety^48^. Under baseline conditions, this hydrophobic group remains shielded within the folded structure. However, when the target sequence interacts with the loop, the hairpin uncoils and exposes the hydrophobic cholesterol moiety to the aqueous environment, driving the rapid aggregation of the AuNPs (**Figure 6**).

**Figure 6.**
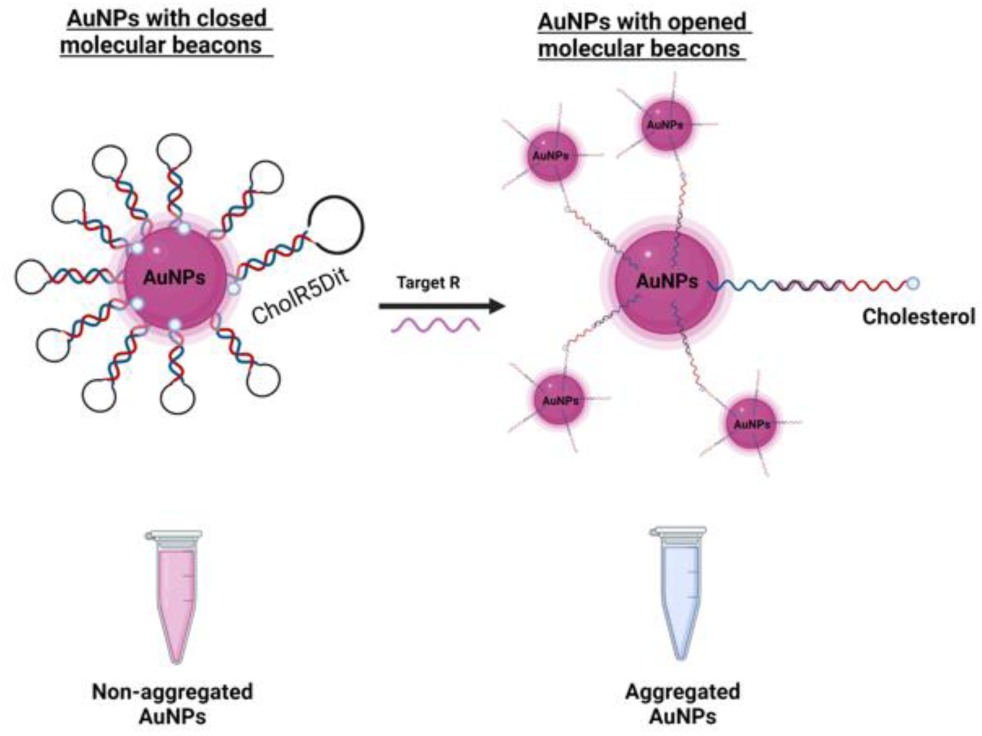
Detection assay of SARS-CoV-2 with AuNPs functionalized with a cholesterol molecular beacon

Building upon our group’s previous adaptation of this strategy for colorimetric SARS-CoV-2 detection using batch-synthesized nanomaterials, we selected the cholesterol-modified molecular beacon targeting the RdRp (R) gene (CholR5Dit, featuring a 5-nucleotide stem) to validate our continuous-flow platform^45^. Before performing the detection assays, the hydrodynamic size and surface charge of the AuNPs functionalized with CholR5Dit under continuous-flow conditions were measured and compared with their batch-produced counterparts. The results revealed that the hydrodynamic diameter and zeta potential of the flow-synthesized AuNPs-CholR5Dit were similar to those prepared under traditional batch conditions. This demonstrated that the AuNPs prepared in the flow reactor exhibit equivalent physicochemical characteristics to the ones produced via batch methods. These results are summarized in **Table 3**.

**Table 3.**
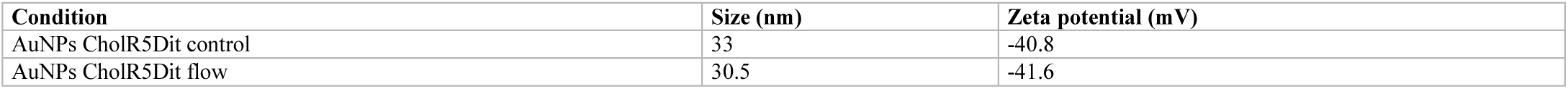
Hydrodynamic diameter and zeta potential of the synthesized AuNPs functionalized with the CholR5Dit probe under batch and continuous-flow conditions.

| **Condition** | **Size (nm)** | **Zeta potential (mV)** |
|---|---|---|
| AuNPs CholR5Dit control | 33 | -40.8 |
| AuNPs CholR5Dit flow | 30.5 | -41.6 |

The results of the detection assay utilizing flow-synthesized AuNPs functionalized with the cholesterol molecular beacon (CholR5Dit) reveal that at initial time (t = 0), AuNPs treated with the specific target (R) and the non-specific control (E) did not exhibit any appreciable colorimetric change upon visual inspection. However, after 60 minutes, a noticeable color change was observed exclusively in the AuNPs treated with the specific target, becoming significantly more intense at 120 minutes (**Figure 7**). This behavior is driven by the specific interaction of target R with the loop of the hairpin design, which induces a change in the probe conformation. This event exposes the cholesterol moieties of the CholR5Dit sequence, previously shielded near the AuNP surface, to the aqueous solvent. Consequently, the hydrophobic cholesterol groups from neighboring AuNPs interact with one another, driving interparticle aggregation. This phenomenon eventually leads to the aggregation and precipitation of the AuNPs at the bottom of the tubes, which is clearly visible at 120 minutes. Conversely, incubation of the AuNPs with the non-specific target E produced no appreciable change in color. These results correlate with a red-shift of the localized surface plasmon resonance (LSPR) peak toward longer wavelengths, which was further confirmed by the UV-Vis absorbance spectra of the samples at 60 and 120 minutes in comparison to the untreated AuNP control (**Figure 8**).

**Figure 7.**
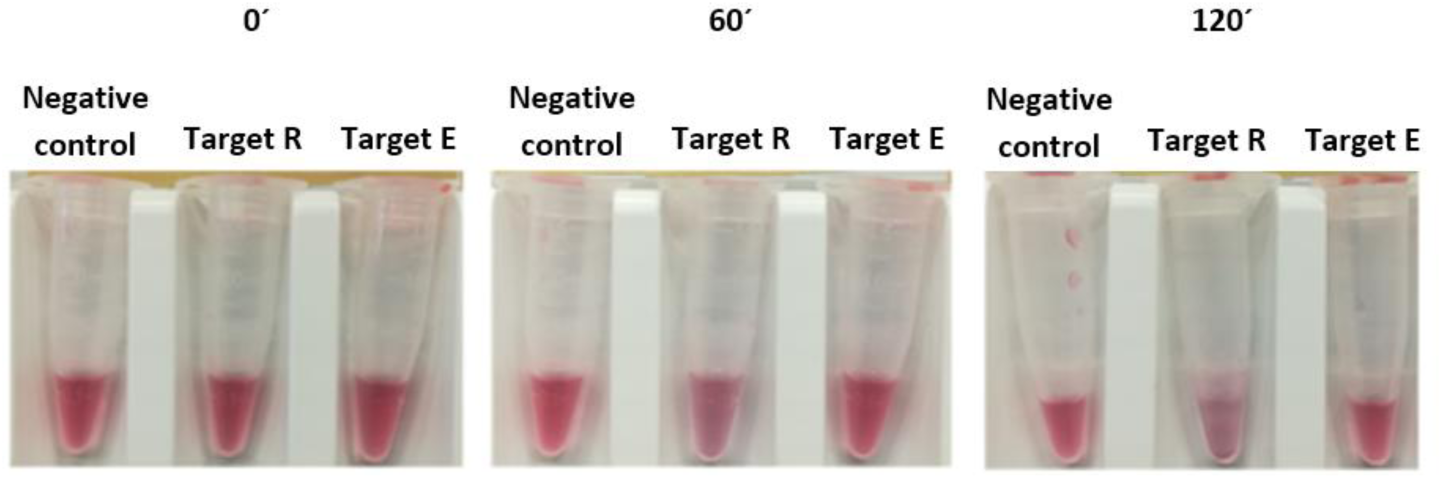
Colorimetric detection assay of the flow-synthesized AuNPs after incubation in the absence of target (negative control), with the specific target (R), and with the non-specific control (E) at t = 0, 60, and 120 minutes.

**Figure 8.**
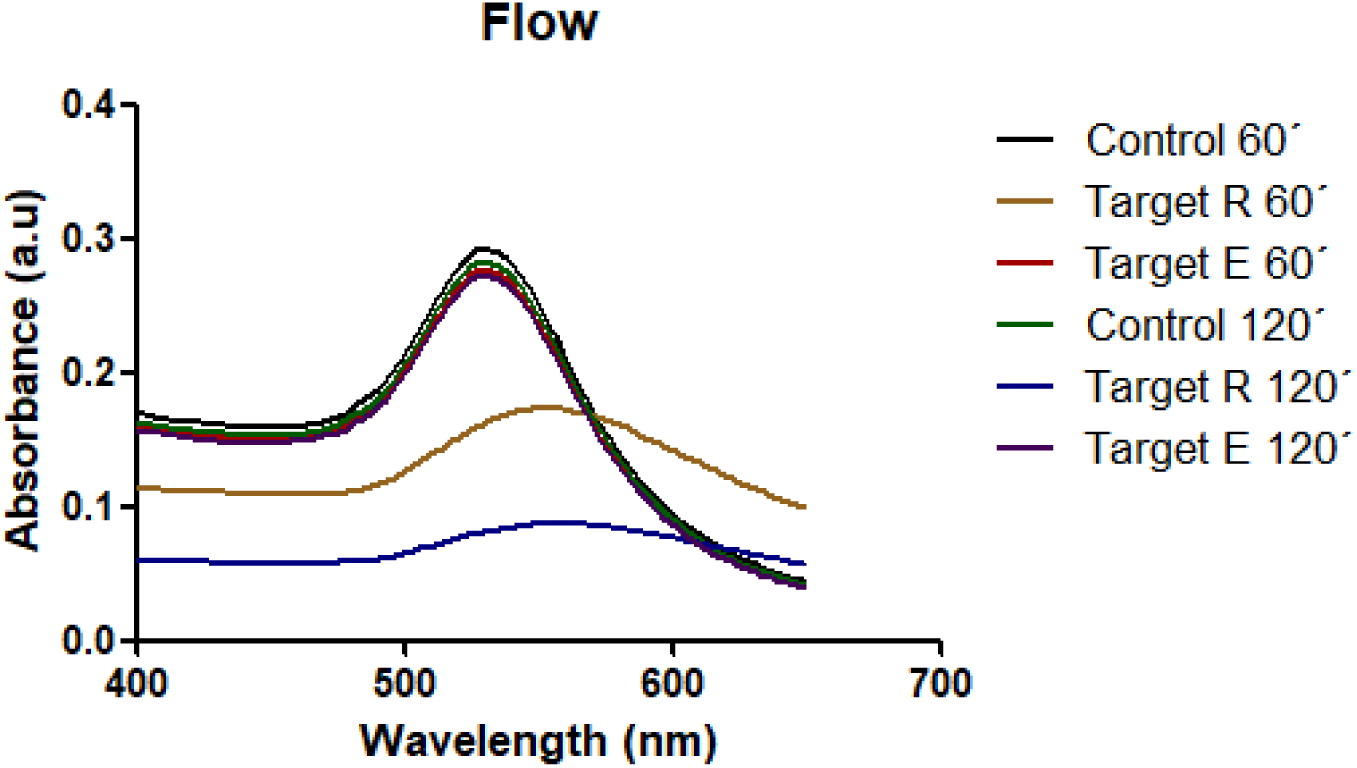
UV-Vis absorbance spectra of the detection assay utilizing flow-synthesized AuNPs after incubation in the absence of target (negative control), with the specific target (R), and with the non-specific control (E) at 60 and 120 minutes.

Crucially, these optical and kinetic results closely mirrored our previously published batch benchmarks. In that earlier study, 20– 25 nm AuNPs functionalized via manual salt-aging with the same 5-nucleotide stem beacon exhibited a distinct naked-eye color shift and a decrease in LSPR absorbance at ∼520 nm within 60 to 120 minutes upon target recognition^45^. The flow-synthesized AuNPs-CholR5Dit reproduced this behavior, with colloidal properties similar to those of the batch-prepared particles (hydrodynamic diameter of 30.5 versus 33 nm and zeta potential of -41.6 versus -40.8 mV, Table 3), and showed a pronounced LSPR red-shift and absorbance drop at 60 and 120 minutes exclusively in the presence of target R, while remaining unreactive toward the non-specific E gene control. This confirms that continuous-flow functionalization reproduces the response time, specificity, and visual readout of the batch sensor without requiring time-consuming salt-aging steps.

#### Detection assay of SARS-CoV-2 with AuNPs functionalized with the biotin molecular beacon

To further expand the diagnostic scope of the molecular beacon architecture previously optimized by our group in batch solutions, we adapted the system to a paper-based lateral flow assay (LFA) format by replacing the hydrophobic cholesterol moiety at the distal terminus with a biotin tag (BiotinR5Dit). A related architecture, in which AuNPs carrying biotin-modified hairpin probes are captured on streptavidin strips, has been reported by our group for the detection of miRNAs^49^. Under this design, the AuNPs bearing the closed, self-hybridized molecular beacon are deposited onto the conjugate pad and subsequently exposed to the specific target. Upon target recognition, the molecular beacon undergoes a conformational change to an open state, thereby exposing the previously masked biotin molecules. These now-accessible biotin moieties are then capable of interacting with the streptavidin immobilized on the test line. Consequently, in this study, AuNPs functionalized with this biotinylated molecular beacon were successfully synthesized using the flow reactor and validated for the detection of the SARS-CoV-2 target (**Figure 9**).

**Figure 9.**
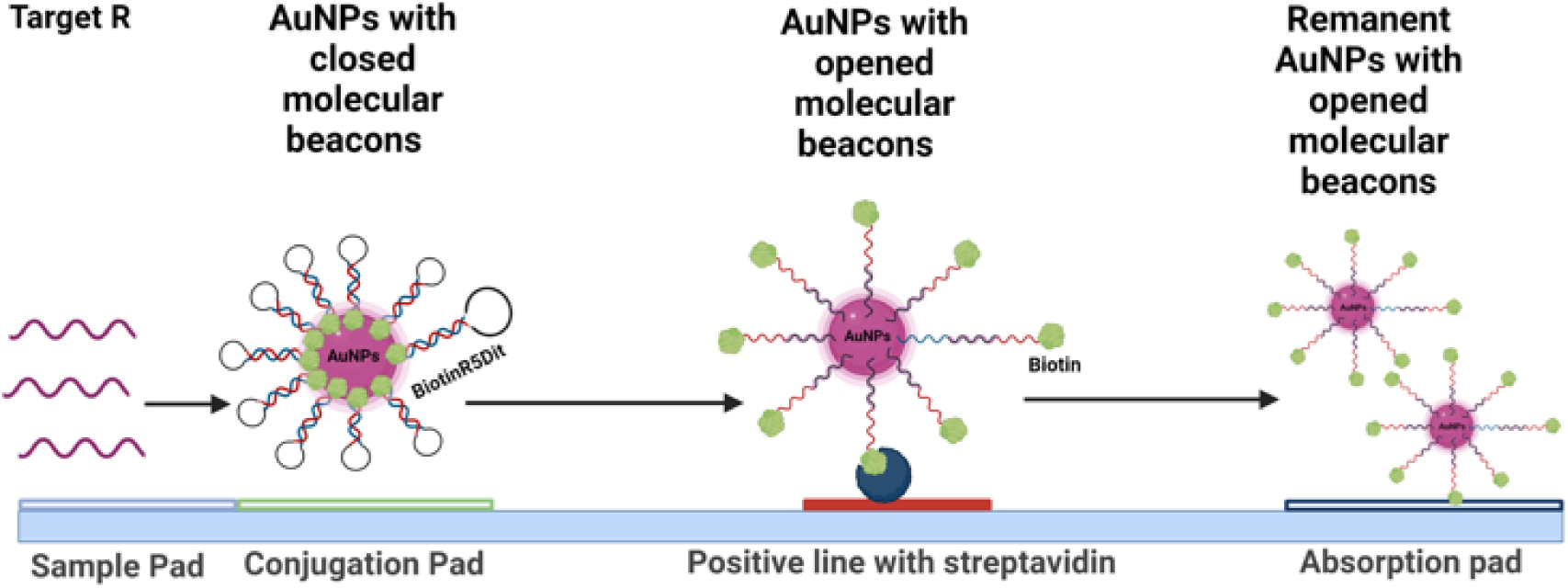
Detection assay of AuNPs functionalized with the biotin molecular beacon.

Prior to the analytical evaluation, the hydrodynamic size and surface charge of the AuNPs functionalized with BiotinR5Dit in the flow reactor were characterized and compared with their batch-synthesized counterparts. The results revealed that both the hydrodynamic diameter and zeta potential of the BiotinR5Dit-AuNPs produced via continuous flow were highly comparable to those obtained under conventional batch conditions. This strong agreement indicates that the microfluidic flow synthesis preserves the core physicochemical attributes and surface characteristics of the nanomaterial. These comparative data are summarized in Table 4.

**Table 4.** Hydrodynamic diameter and zeta potential of the synthesized AuNPs functionalized with the BiotinR5Dit probe under batch and continuous-flow conditions.

| **Entry** | **Condition** | **Size (nm)** | **Zeta potential (mV)** |
|---|---|---|---|
| **1** | AuNPs BiotinR5Dit control | 28.93 | -38.5 |
| **2** | AuNPs BiotinR5Dit flow | 25.13 | -37.7 |

The functional evaluation of the continuous-flow synthesized AuNPs functionalized with the biotinylated molecular beacon (BiotinR5Dit) revealed the successful formation of distinct red lines across all tested conditions upon incubation with target R. This biosensing process operated in a concentration-dependent manner, where a decrease in target concentration correlated with a reduction in test line intensity, establishing a limit of detection (LOD) at 100 nM. For these assays, the functionalized AuNPs were integrated into the conjugate pad and evaluated against target R concentrations of 1000, 500, 250, 100, and 25 nM (**Figure 10 and Figure 11**). Mechanistically, the introduction of the specific target R triggers the opening of the molecular beacon’s hairpin structure, exposing the biotin moiety for subsequent affinity capture by the streptavidin immobilized at the test line.

**Figure 10.**
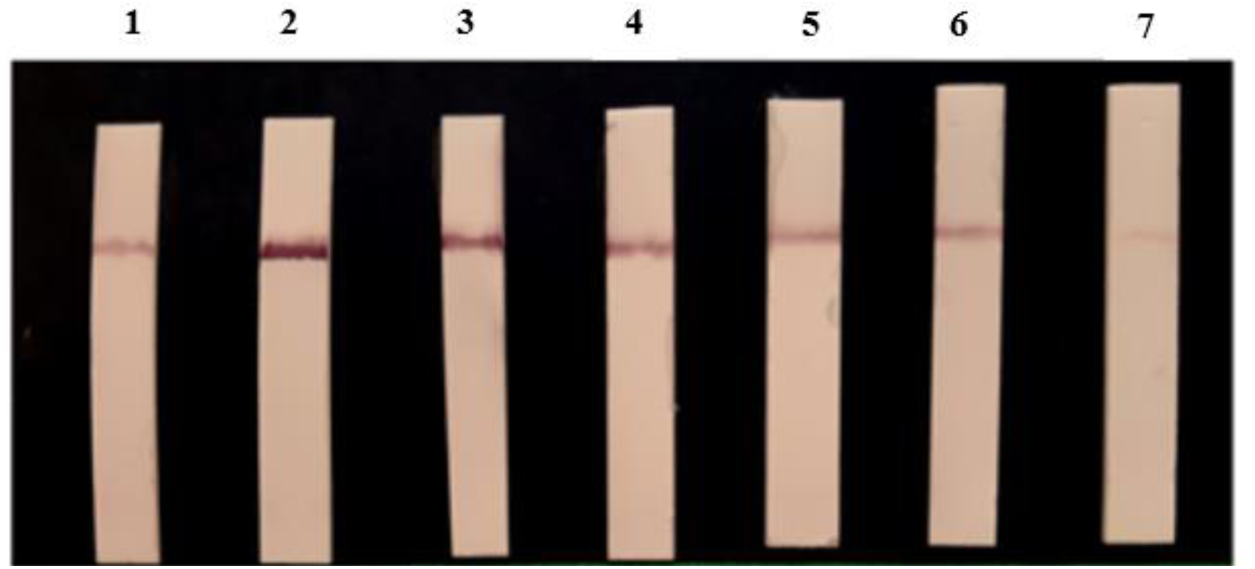
Evaluation of SARS-CoV-2 target R detection using BiotinR5Dit-functionalized AuNPs. Lateral flow strips after incubation under different experimental conditions: (1) negative control (without target); specific target R at final concentrations of (2) 1000 nM, (3) 500 nM, (4) 250 nM, (5) 100 nM, and (6) 25 nM; and (7) a PEGylated AuNP control.

**Figure 11.**
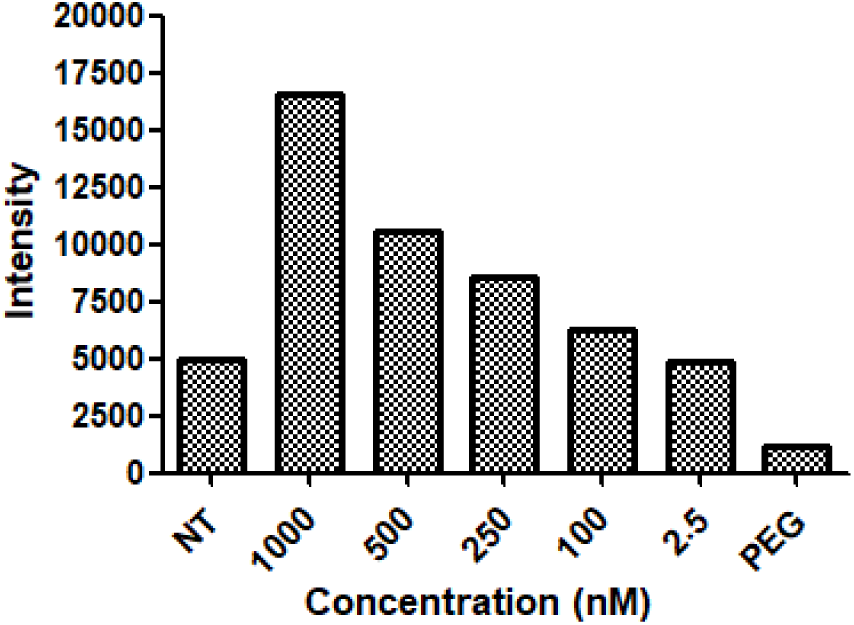
Quantitative evaluation of Ponceau S-treated lateral flow assay profiles. Signal intensities of the functionalized AuNPs recorded after protein staining under the following conditions: (1) negative control, (2–6) specific target R at decreasing concentrations (1000, 500, 250, 100, and 25 nM), and (7) PEGylated AuNP control.

Conversely, a clear signal was observed in the negative control, which could be attributed to a fraction of BiotinR5Dit molecules failing to adopt the proper closed hairpin conformation on the gold surface, or to non-specific interactions between the AuNP core and streptavidin. Furthermore, the subtle signal line detected in the AuNPs-PEG control suggests an insufficient PEG surface density to completely suppress these non-specific matrix interactions.

#### Detection assay of SARS-CoV-2 with AuNPs functionalized with RNA by CRISPR-Cas13a

CRISPR-Cas enzymes have been widely utilized as point-of-care testing (POCT) platforms for the sensitive and specific detection of nucleic acids^50^. Among these approaches, CRISPR-Cas colorimetric sensors are particularly appealing due to their simplicity, ease of use, and low cost. In this context, López-Valls et al. developed CASCADE (CRISPR/CAS-based Colorimetric nucleic Acid DEtection), a SARS-CoV-2 sensor wherein the CRISPR-Cas13 ribonucleoprotein (RNP) complex is activated upon recognition of the target RNA, which in turn triggers its non-specific collateral trans-cleavage activity^46^. This promiscuous activity was exploited to cleave the RNA oligonucleotides that stabilized the AuNPs. Consequently, the degradation of the protective RNA shell induced interparticle aggregation, enabling visual detection by the naked eye. In that earlier work using batch-synthesized nanomaterials, 12 nm AuNPs coated via manual salt-aging with a 33-nucleotide ssRNA probe (RNA3) enabled the naked-eye detection of 25 nM

SARS-CoV-2 Orf1ab target RNA within 15–30 minutes without requiring centrifugation or salt addition. Building upon this framework, we produced the equivalent 13 nm AuNPs functionalized with the same 33-nt sequence (CRCS1Dit) entirely under continuous-flow conditions and subsequently evaluated them for the detection of SARS-CoV-2 viral targets via a CRISPR-Cas13a assay (**Figure 12**).

**Figure 12.**
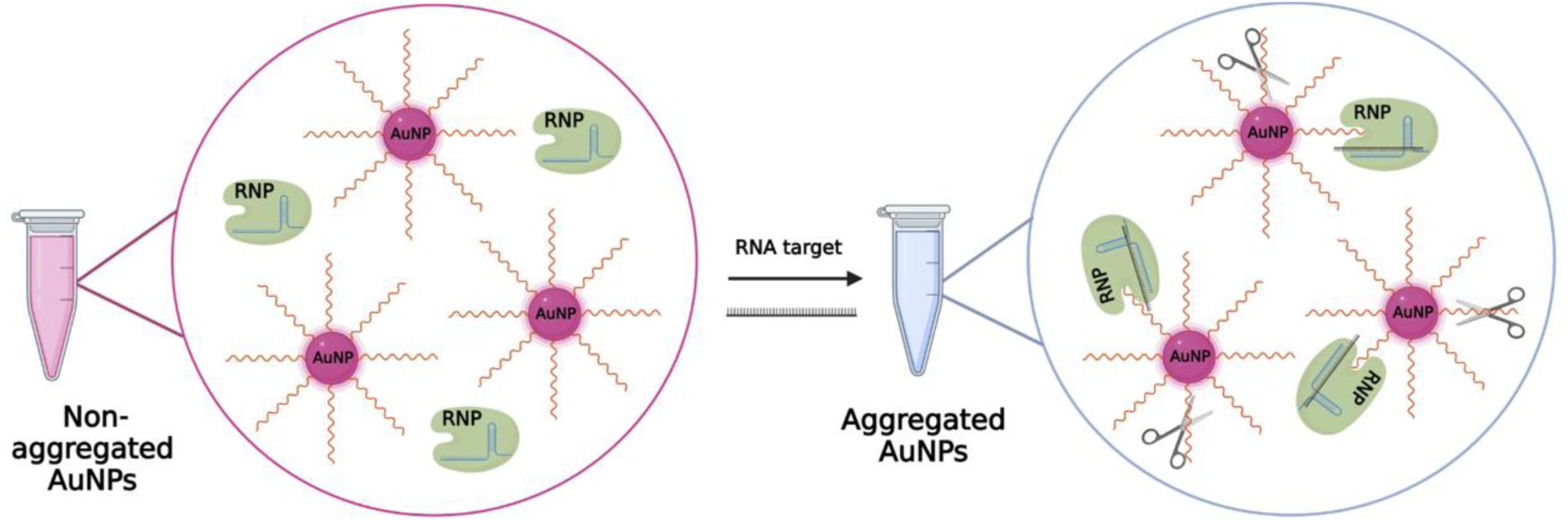
Detection assay of SARS-CoV-2 with AuNPs functionalized with RNA by CRISPR-Cas13a

Before performing the detection assays, the hydrodynamic size and surface charge of the AuNPs functionalized with CRCS1Dit under continuous-flow conditions were measured and compared with their batch-produced counterparts. The results revealed that the hydrodynamic diameter and zeta potential of the flow-synthesized AuNPs-CRCS1Dit were similar to those prepared under traditional batch conditions. This demonstrated that the AuNPs prepared in the flow reactor exhibit equivalent physicochemical characteristics to the ones produced via batch methods. These results are summarized in **Table 5**.

**Table 5.** Hydrodynamic diameter and zeta potential of the synthesized AuNPs functionalized with the CRCS1Dit probe under batch and continuous-flow conditions.

| **Condition** | **Size (nm)** | **Zeta potential (mV)** |
|---|---|---|
| AuNPs CRCS1Dit control | 35.14 | -53.4 |
| AuNPs CRCS1Dit flow | 40.97 | -52.0 |

The results of the CRISPR-Cas13a detection assay utilizing flow-synthesized AuNPs functionalized with the RNA stabilizing molecule (CRCS1Dit) revealed an immediate colorimetric transition from intense red to a lighter red hue upon addition of target R. This aggregation process further progressed at 15 minutes (turning purple) and 30 minutes (turning gray). Finally, a distinct pellet was formed at 60 minutes. In contrast, no colorimetric changes were observed in the absence of the target (RNP control), and the blank condition containing neither the protein nor the target also remained stable (**Figure 13**). In this sensing mechanism, the interparticle aggregation is not driven by hydrophobic interactions but rather by the target-activated non-specific collateral trans-cleavage activity of the ribonucleoprotein (RNP) complex, which rapidly cleaves the surrounding RNA strands. Specifically, the enzymatic degradation of the CRCS1Dit sequence, which acts as the electrostatically stabilizing shell, disrupts the steric and electrostatic barriers of the nanomaterial. Crucially, the observation that the RNP complex alone (in the absence of target R) did not induce AuNP aggregation confirms that the ribonucleoprotein framework does not compromise the basal colloidal stability of the AuNPs.

**Figure 13.**
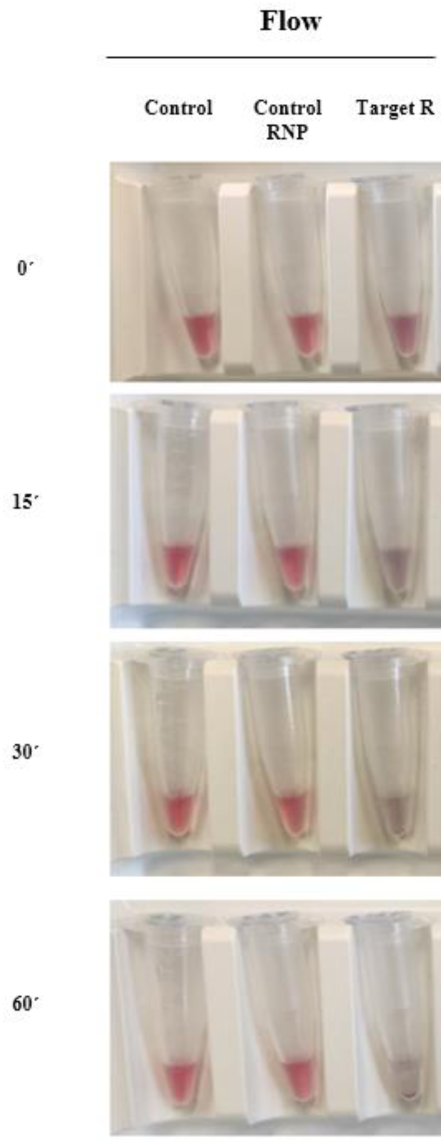
Colorimetric CRISPR-Cas13a detection assay of the flow-synthesized AuNPs after incubation in the absence of both the target and the RNP complex (blank control), with the RNP alone (RNP control), and with the specific target (R) at a concentration of 25 nM at t = 0, 15, 30, and 60 minutes.

A direct comparison between the flow-produced nanoconjugates and our previously published batch CASCADE sensor shows comparable performance with both fabrication methods. In the original batch study, 25 nM of target was detected by the naked eye within 15–30 minutes; in the flow-prepared sample, the suspension challenged with 25 nM target R changed from red to purple at 15 minutes and to gray at 30 minutes, with precipitation at 60 minutes, while the RNP and blank controls remained stable. These findings indicate that the flow-prepared AuNPs reproduce the detection time and naked-eye readout of the manual CASCADE system.

### Therapeutic applications: Targeted drug delivery and gene silencing

Having validated the diagnostic reliability of the flow-synthesized nanoconjugates against established batch benchmarks, we next evaluated the versatility of our modular platform for therapeutic applications. To demonstrate that the reactor can handle chemically distinct bioactive payloads, we selected two examples based on two types of therapeutic agents: (i) a chemotherapeutic drug, SN38, combined with the nucleolin-targeting aptamer AS1411 for tumor targeting, and (ii) a nucleic acid, the antisense oligonucleotide ASO34a, for gene silencing.

### Cell viability assay with AuNPs functionalized with nucleic acids and SN38

Traditional cancer chemotherapies have been hampered by severe limitations, such as significant systemic toxicity due to poor bioavailability, metabolic instability caused by in vivo degradation, inherent low solubility, and P-glycoprotein (P-gp)-mediated multidrug resistance^51^. SN38, the active metabolite of irinotecan, is a potent topoisomerase I inhibitor with proven efficacy against colorectal, lung, and breast cancers, among others; however, its clinical translation is severely impeded by these exact physicochemical and pharmacological drawbacks^52^. To overcome these hurdles, AuNPs have emerged as promising drug carriers due to their high surface area-to-volume ratio, which provides abundant binding sites for drug loading, along with enhanced drug solubility, facile surface functionalization, and excellent biocompatibility. On the other hand, the aptamer AS1411, a 26-base G-rich oligodeoxynucleotide sequence with intrinsic apoptosis-inducing activity, has shown promising clinical activity and is being widely utilized as a tumor-targeting agent due to the specific overexpression of nucleolin on the surface of cancer cells^53^. Consequently, in the present study, AuNPs co-functionalized with SN38 and AS1411 molecules were successfully engineered utilizing the continuous-flow reactor and subsequently evaluated in cell viability assays (**Figure 14**).

**Figure 14.**
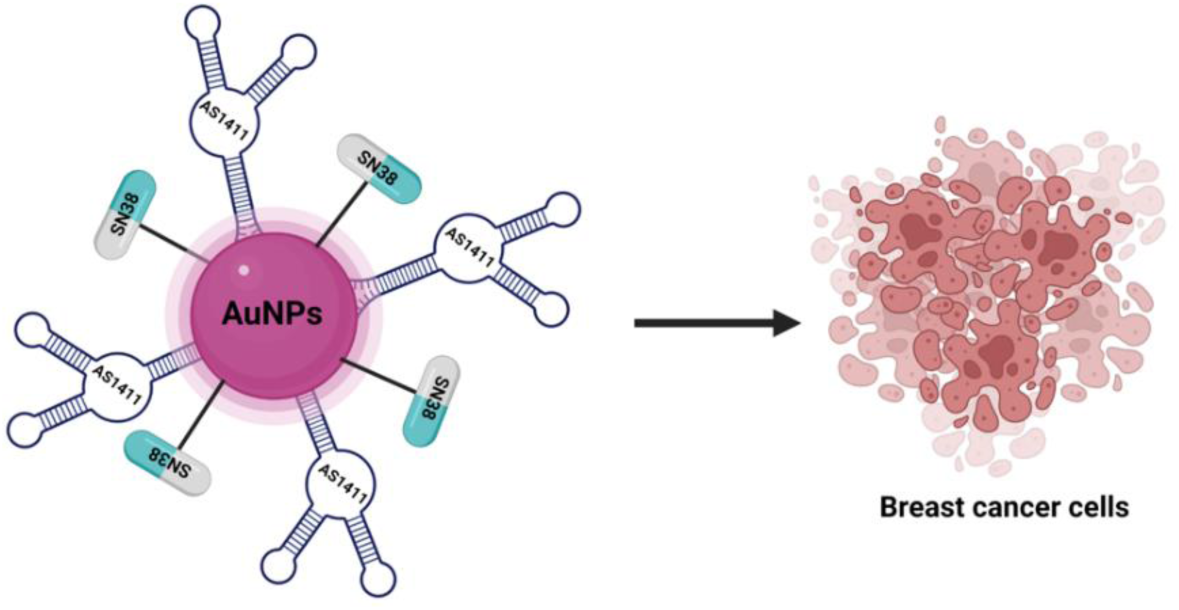
Cell viability assay with AuNPs functionalized with nucleic acids and SN38.

Prior to the cell viability evaluations, the hydrodynamic size and surface charge of the AuNPs functionalized with PEG, SN38Dit + AS1411Dit, SN38Dit + PEGDit, and SN38Dit + SCRDit under continuous-flow conditions were measured and compared with their batch-produced counterparts. The hydrodynamic diameters of the flow-synthesized AuNPs were comparable to those of the batch-prepared ones, as were the zeta potentials of the formulations containing nucleic acids. In contrast, the PEG-containing formulations showed a less negative zeta potential in flow (-13.6 versus -33.5 mV for AuNPs PEG, and -14.3 versus -22 mV for SN38Dit + PEGDit), which may reflect a higher PEG loading under flow conditions. These results are summarized in **Table 6**.

**Table 6.** Hydrodynamic diameter and zeta potential data of the synthesized AuNPs functionalized with PEG, SN38Dit + Perdita, SN38Dit + SCRDit, and SN38Dit + AS1411Dit under batch and continuous-flow conditions.

| **Condition** | **Size (nm)** | **Zeta potential (mV)** |
|---|---|---|
| AuNPs PEG control | 46.8 | -33.5 |
| AuNPs PEG flow | 48 | -13.6 |
| AuNPs SN38Dit + PEGDit control | 23.6 | -22 |
| AuNPs SN38Dit + PEGDit flow | 26.9 | -14.3 |
| AuNPs SN38Dit + SCRDit control | 37.1 | - 45 |
| AuNPs SN38Dit + SCRDit flow | 41.7 | -42.3 |
| AuNPs AS1411Dit + SN38Dit control | 42.6 | -43.6 |
| AuNPs AS1411Dit + SN38Dit flow | 38.5 | -41.1 |

The results of the cell viability assays evaluating the flow-synthesized AuNPs functionalized with SN38Dit + AS1411Dit, SN38Dit + PEGDit, and SN38Dit + SCRDit in MDA-MB-231 breast cancer cells, across various incubation times (24, 48, and 72 h) and drug concentrations (0.5, 1, 3, 8, and 15 µM), demonstrated that control AuNPs without SN38 exhibited negligible cytotoxicity, with the PEG-functionalized formulation demonstrating the highest biocompatibility. Conversely, all AuNPs loaded with SN38 induced significant cytotoxicity. A comparable cytotoxic profile was observed between the AuNPs + PEGDit and AuNPs + SCRDit treatment groups. Notably, the most pronounced therapeutic efficacy was achieved by the nanoformulation co-loaded with SN38 and the AS1411 aptamer (**Figure 15 and Figure 16**). This enhanced anti-proliferative effect can be attributed to the active targeting role of the AS1411Dit sequence, which has been reported to facilitate receptor-mediated cellular internalization and trigger intrinsic apoptotic pathways.

**Figure 15.**
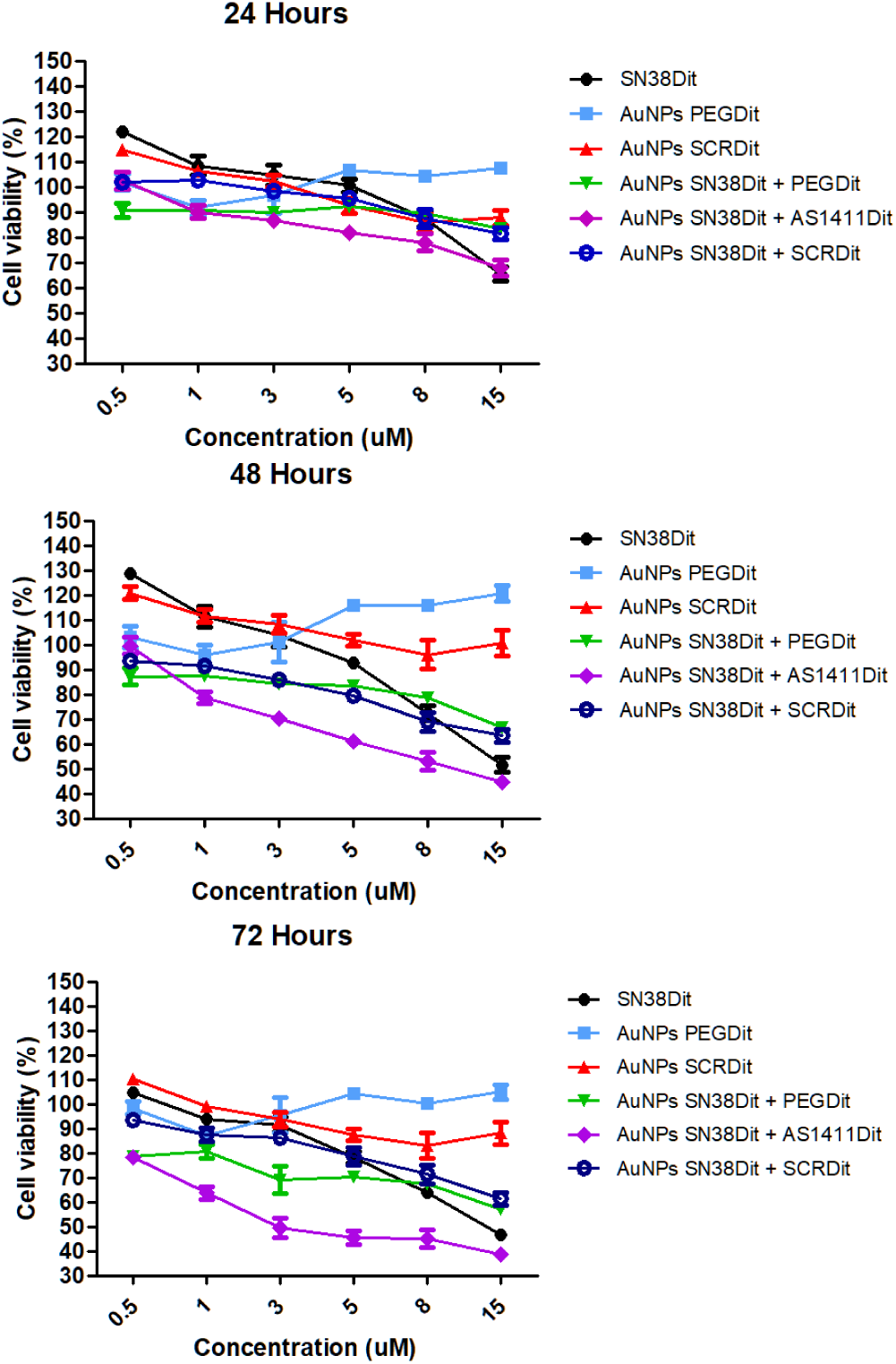
In vitro cell viability assays of MDA-MB-231 breast cancer cells treated with flow-synthesized AuNPs functionalized with PEG, SN38Dit + PEGDit, SN38Dit + SCRDit, and SN38Dit + AS1411Dit after 24, 48, and 72 hours of incubation across a concentration range from 0.5 to 15 µM.

**Figure 16.**
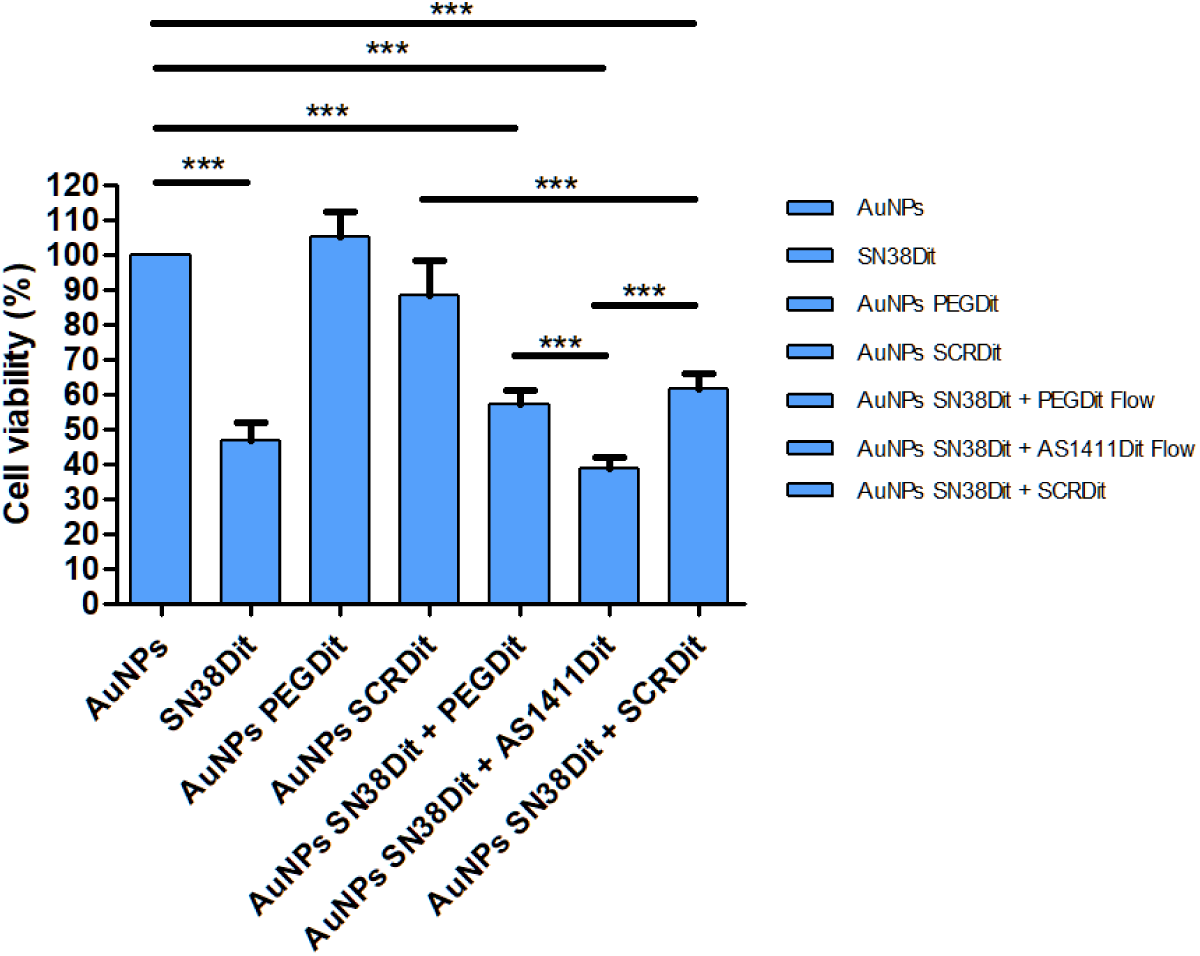
In vitro cell viability of MDA-MB-231 breast cancer cells after 72 hours of incubation with flow-synthesized AuNPs functionalized with PEG, SN38Dit + PEGDit, SN38Dit + SCRDit, and SN38Dit + AS1411Dit at a fixed concentration of 15 μM.

### Gene silencing assay with AuNPs functionalized with ASO34a mimic

Antisense oligonucleotides (ASOs) are synthetic, single-stranded nucleic acid sequences designed to hybridize with high specificity to target complementary RNA sequences through Watson-Crick base pairing. Upon binding, ASOs modulate gene expression primarily by triggering RNase H-mediated mRNA degradation or sterically blocking translation, thereby achieving a precise gene silencing effect at the post-transcriptional level^54^. This therapeutic strategy allows for the modulation of key signaling pathways involved not only in tumorigenesis (such as cell proliferation, metastasis, and apoptosis evasion) but also in the pathogenesis of numerous rare and genetic diseases. However, naked ASOs suffer from rapid enzymatic degradation and poor intracellular uptake^55^. To overcome these delivery hurdles, AuNPs have emerged as highly efficient platforms for the surface immobilization and delivery of ASOs, significantly enhancing their chemical biostability against nuclease degradation and promoting receptor-mediated cellular internalization^56^. Based on this, the AuNPs synthesized within the continuous-flow reactor were functionalized with the antisense oligonucleotide ASO34a and subsequently evaluated in a gene silencing assay (**Figure 17**).

**Figure 17.**
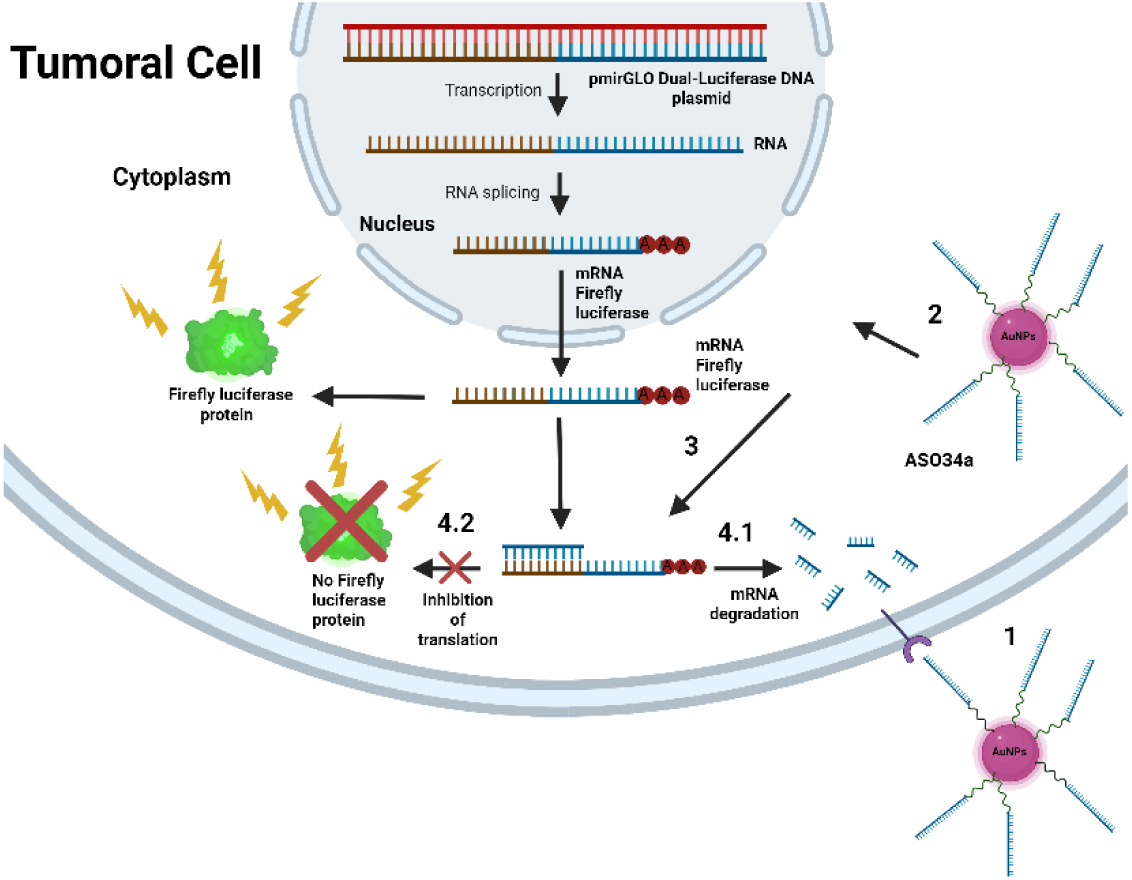
Silencing assay of AuNPs functionalized with ASO34a mimic

Following functionalization, the hydrodynamic size and surface charge of the AuNPs functionalized with the ASO34a mimic sequence within the continuous-flow reactor were measured and compared with their batch-produced counterparts. The hydrodynamic diameters were comparable (36 nm for batch and 33.6 nm for flow), whereas the flow-synthesized conjugate showed a more negative zeta potential (-45.4 versus -37.9 mV). These results are summarized in **Table 7**.

**Table 7.** Hydrodynamic diameter and zeta potential data of the synthesized AuNPs functionalized with thiolated ASO34a mimic under batch and continuous-flow conditions.

| **Condition** | **Size (nm)** | **Zeta potential (mV)** |
|---|---|---|
| AuNPs ASO34a mimic thiol control | 36 | -37.9 |
| AuNPs ASO34a mimic thiol flow | 33.6 | -45.4 |

Luciferase assays were performed to evaluate the silencing efficiency of flow-synthesized AuNPs functionalized with ASO34a at final concentrations of 3, 6, and 9 nM in MDA-MB-231 and PANC-1 cell lines after 24 hours of incubation. The results confirmed a significant knockdown effect across both cell models. Specifically, in the breast cancer cell line (MDA-MB-231), luciferase expression was reduced to 69.0%, 66.0%, and 52.8%, respectively. In the pancreatic cancer cell line (PANC-1), the remaining expression levels were 66.6%, 70.0%, and 69.0% for the corresponding concentrations (**Figure 18**), with no dose dependence over the range tested.

**Figure 18.**
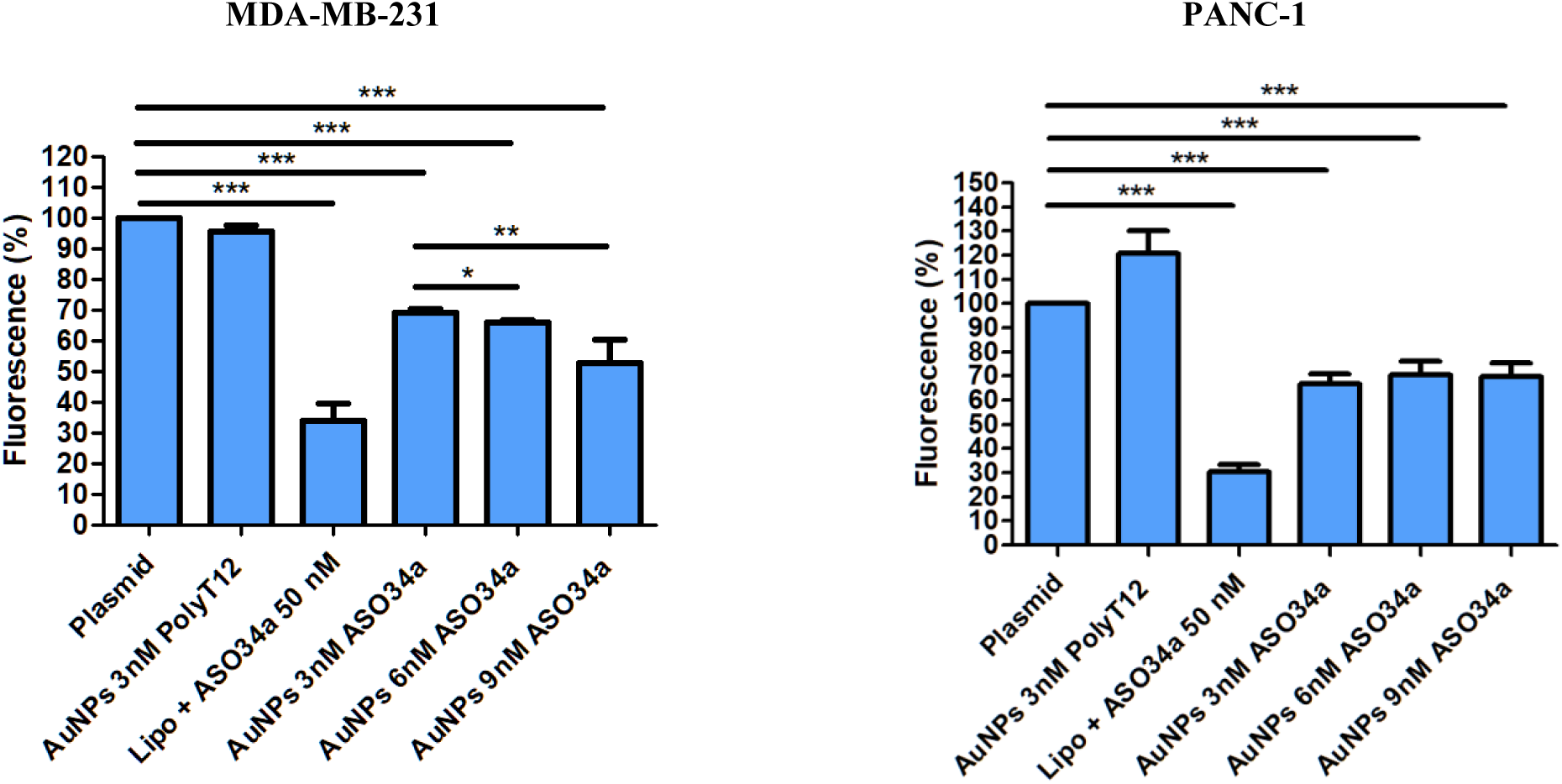
In vitro gene silencing efficiency. **Relative luciferase expression** of MDA-MB-231 (breast cancer) and PANC-1 (pancreatic cancer) cell lines 24 h post-treatment with flow-synthesized AuNPs functionalized with ASO34a mimic at final concentrations of 3, 6, and 9 nM. Untreated plasmid and AuNPs functionalized with PolyT12 were used as negative controls, while Lipofectamine (Lipo) served as the positive transfection control. Cell viability under the same conditions is shown in Supplementary Figure 1.

## Discussion

The landscape of gold-based nanomaterials that have reached the market as medical devices, principally through lateral flow assays, or are currently undergoing clinical trials, reveals that manufacturing pressures center not solely on the gold core itself, but also on the high-value molecules bound to its surface^3^. From an academic perspective, the functionalization of AuNPs can encompass even more complex designs, with multi-layered architectures of molecules of entirely different chemical natures, including hydrophilic polymers, highly charged oligonucleotides, hydrophobic chemotherapeutics, and targeting proteins or aptamers^57^.

Traditionally, the synthesis of these functionalized gold nanoparticles has been performed in batch reactors, which suffer from poor mass/heat transfer leading to poor control over grafting densities, and low batch-to-batch reproducibility. In this context, continuous-flow reactors have emerged as a highly attractive alternative, owing to their superior mass- and heat-transfer capabilities^6^. These enhanced fluidic properties offer a robust paradigm shift to streamline the fabrication of multi-component nanoconstructs, thereby paving the way for their successful translation into commercial and clinical realities. Currently, there is a plethora of works about the synthesis of gold nanoparticles of different nature in flow reactors^58^. More recently, the focus has turned to reproducibility studies and even modification of AuNPs with polyethylene glycol (PEG) architectures^27,28^. However, the stable and controlled functionalization of these platforms with more complex or multi-layered molecular targets remains a challenge that hinders their translation.

To solve this, we have developed a modular, continuous-flow millifluidic platform capable of performing both the automated synthesis of citrate-stabilized AuNPs and their subsequent multi-step covalent functionalization in a single, streamlined process. Specifically, we have designed two distinct fluidic reactor configurations to adapt Turkevich-synthesized AuNPs to two well-established protocols for oligonucleotide surface modification: the conventional salt-aging strategy and the low-pH-driven adsorption methodology. The mechanism of the salt-aging protocol is based on the mitigation of electrostatic barriers via controlled electrolyte addition, allowing densely charged ligands like oligonucleotides to approach and covalently bind to the AuNP surface. The primary limitation of this method resides in its operational timeline. Because rapid salt addition triggers immediate colloidal collapse, the aging process requires a meticulous, hours-long stepwise increase in salt concentration. In typical laboratory workflows, this translates into tedious incubation periods that regularly span from 16 to over 24 hours^59^. Such extended time requirements represent a major barrier to scalability even in flow reactors. In this work, the utilization of dithiolane linkers (derived from lipoic acid modifications) proved critical for achieving fast and robust covalent attachment within the short residence times (approximately 20 minutes) in the reactor. While the utilization of dithiolane anchors substantially increases the thermodynamic stability of the Au–S bonds, offering an optimal configuration for robust biosensing platforms, therapeutic applications often demand an opposing mechanism. For medical purposes, specific molecular payloads must be efficiently released from the nanoparticle surface to exert their intended therapeutic efficacy at the target site. While it is possible to design sophisticated dithiolane architectures equipped with cleavable release mechanisms, such platforms remain considerably more expensive and complex to develop compared to simpler alternatives^60^. To overcome the kinetic limitations of conventional protocols, the low-pH citrate buffer method has emerged as a salt-free alternative for the ultra-fast functionalization of gold nanoparticles with thiol-modified oligonucleotides. This strategy relies on shifting the local environment to an acidic range (typically using a citrate buffer at pH 3.0) to modulate the ionization state of the nucleic acid bases. At this low pH, the adenine and cytosine bases undergo partial protonation, which effectively neutralizes the net negative charge of the oligonucleotide backbone and screens the electrostatic repulsion between the macromolecule and the citrate-stabilized AuNP surface^59^. Consequently, the hydrophobic and van der Waals attractive forces dominate, driving an almost instantaneous, high-density adsorption of the cargo onto the gold facets. Remarkably, while batch salt-aging requires 16 to over 24 hours, the low-pH-driven mechanism achieves optimal grafting densities within seconds to a few minutes (typically under 1 to 5 minutes) without inducing colloidal collapse, offering an ideal framework for the modification of AuNPs with interference oligonucleotides. Indeed, traditionally, therapeutic gold nanoparticles designed for gene regulation in cancer have relied on oligonucleotides functionalized with simple monothiol groups (-SH) to achieve surface immobilization^61^. These monothiol-based configurations are highly susceptible to premature degradation in the tumor microenvironment due to competitive displacement by intracellular glutathione (GSH), which is significantly overexpressed in cancer cells to counteract the high levels of reactive oxygen species (ROS) generated by their accelerated metabolism. In this work, the ASO34a was attached to the AuNPs through a single thiol group using the low-pH module, an arrangement in which the oligonucleotide can be displaced by intracellular glutathione, while avoiding the long salt-aging step^62^.

The potential of flow reactors is evidenced by the clear correlation between the physicochemical properties (hydrodynamic diameters and, except for the PEG-containing formulations, surface charge) of the flow-synthesized nanoconjugates and their batch counterparts. Furthermore, functional validation assays performed across both setups demonstrated that the flow-produced hybrid AuNPs retained their biological and diagnostic activity, executing their intended mechanisms with high efficiency across the different designs. For the two diagnostic assays previously developed in batch (cholesterol molecular beacon and CASCADE), the flow-prepared AuNPs reproduced the specificity, response time, and naked-eye readout reported for the batch-prepared nanoparticles. The lateral flow format and the therapeutic systems were evaluated here with flow-prepared AuNPs and compared with batch-prepared conjugates only at the physicochemical level (Tables 4 to 7). For the optical, colorimetric nucleic acid sensors (both the hydrophobic cholesterol-driven and the CRISPR-Cas13a-mediated platforms), the lateral flow assay sensor, and targeted drug delivery (SN38 + AS1411) the flow-modified AuNPs performed well without requiring the conventional, tedious "salt-aging" protocol. A similar trend was observed for the AuNPs modified with the ASO34a via the low-pH method, confirming that the continuous-flow platform can successfully be adapted to this methodology.

In detail, the solution-based assays, the cholesterol molecular beacon (CholR5Dit) operated through targeted hairpin loop opening, exposing the hydrophobic core to the aqueous phase and triggering visible interparticle aggregation and LSPR red-shifts. Similarly, the CRISPR-Cas13a biosensor leveraged target-activated trans-cleavage kinetics to enzymatically strip away the protective CRCS1Dit stabilizing ribonucleic shell, breaking down electrostatic and steric barriers to induce a rapid, naked-eye color change from red to purple/gray. When translated to paper-based lateral flow assays (LFAs) using the biotinylated beacon (BiotinR5Dit), the system successfully achieved a limit of detection (LOD) of 100 nM for target R in a concentration-dependent manner. However, the appearance of background signals in the negative and AuNP-PEG controls warrants careful biochemical examination. The visible line in the negative control (absence of target) points toward a small subpopulation of BiotinR5Dit probes failing to correctly adopt the closed hairpin conformation upon surface immobilization, an effect also described for AuNPs carrying biotin-modified hairpin probes at high surface density^49^, thereby leaving the biotin molecule constitutively exposed for streptavidin affinity capture. Alternatively, this background could stem from direct, non-specific interactions between unpassivated regions of the gold surface and the immobilized streptavidin matrix. This second hypothesis is strongly supported by the subtle line observed in the AuNPs-PEG control, which implies that the PEG packing density achieved under these specific flow configurations was close to, but slightly below, the absolute threshold required to fully eliminate non-specific macromolecular adsorption. Further optimization, such as adjusting the probe and PEG densities on the AuNP surface, is therefore warranted.

From a therapeutic perspective, the co-functionalized nanoconjugates demonstrated outstanding performance. In the drug delivery modules, the simultaneous conjugation of the topoisomerase I inhibitor SN38 and the nucleolin-targeting AS1411 aptamer yielded a formulation with superior anti-proliferative activity against triple-negative breast cancer cells (MDA-MB-231) compared to non-targeted controls. This enhanced cytotoxicity may be attributed to the receptor-mediated endocytosis driven by the AS1411 ligand, which targets the overexpressed nucleolin on the cancer cell surface and facilitates intracellular accumulation of the cytotoxic cargo. Furthermore, the immobilization of the antisense oligonucleotide ASO34a onto the AuNP surface yielded a potent post-transcriptional gene silencing platform. This system achieved significant luciferase knockdown in both breast (MDA-MB-231) and pancreatic (PANC-1) cancer cell lines, achieving a minimum of 30% silencing efficiency within 24 hours of exposure. Crucially, the nanoconjugates exhibited an excellent biocompatibility profile, displaying no significant cytotoxic effects in either of the evaluated cell lines at the therapeutic doses (**Supplementary Figure 1**).

Taken together, these findings demonstrate that continuous-flow chemistry is not merely an alternative manufacturing route, but an enabling, highly versatile technology for nanomedicine. By integrating core synthesis and diverse multi-component functionalization within a single automated pipeline, this approach demonstrates that complex modifications of AuNPs can be successfully performed in flow reactors, thereby narrowing the gap between laboratory discovery and clinical manufacturing.

## Conclusions

In this work, we have successfully developed a versatile and modular continuous-flow millifluidic platform that integrates both the automated synthesis of citrate-stabilized gold nanoparticles and their subsequent multi-step surface modification within a single, streamlined pipeline. By precisely controlling the reagent stoichiometry, uniform AuNPs of 13 and 22 nm were reliably produced. The flow-derived nanoconjugates showed hydrodynamic sizes comparable to those of the batch formulations, and comparable zeta potentials except for the PEG-containing ones, thereby supporting the reliability of our fluidic platform.

The chemical architecture of the platform was validated through two distinct functionalization strategies tailored for oligonucleotide immobilization. First, the integration of bidentate dithiolane anchors proved critical to achieving rapid and efficient covalent ligand exchange within the short residence times of the fluidic modules. This strategy enabled the generation of high-performance colorimetric biosensors for SARS-CoV-2 (utilizing cholesterol-driven or CRISPR-Cas13a-mediated mechanisms) and paper-based lateral flow assays, completely bypassing the tedious, hours-long "salt-aging" steps standard in benchtop protocols. Second, the platform was successfully adapted to a salt-free, low-pH-driven adsorption methodology, expanding the operational plasticity of automated flow manufacturing.

From a biomedical perspective, the flow-produced nanoconjugates demonstrated diagnostic and therapeutic activity in vitro. In diagnostic applications, replicating our previously published batch-synthesized SARS-CoV-2 assays confirmed that continuous-flow functionalization preserves the target specificity, response time, and naked-eye readout of both the cholesterol molecular beacon and CRISPR-Cas13a (CASCADE) sensors, while enabling seamless adaptation to paper-based lateral flow formats without time-consuming salt-aging steps. In therapeutic applications, the platform successfully accommodated two chemically distinct classes of bioactive payloads: a targeted small-molecule chemotherapeutic formulation co-conjugated with SN38 and the AS1411 aptamer, which exhibited potent anti-proliferative activity against triple-negative breast cancer cells (MDA-MB-231), and a nucleic acid-based system functionalized with the antisense oligonucleotide ASO34a via the low-pH module, which achieved significant post-transcriptional gene silencing (driving a minimum of 30% luciferase knockdown within 24 hours in both breast and pancreatic PANC-1 cancer cell lines). Importantly, the ASO34a-functionalized AuNPs showed no significant cytotoxicity in either cell line at the concentrations used for gene silencing (**Supplementary Figure 1**), and the PEG-functionalized AuNPs lacking SN38 showed negligible toxicity in MDA-MB-231 cells.

In conclusion, this automated millifluidic platform represents an enabling technology that unifies the synthesis and intricate, multi-component functionalization of AuNPs. By successfully overcoming the limitations of standard fluidic systems, this approach effectively bridges the gap between simple surface attachments and highly complex structural modifications under continuous manufacturing workflows.

## Acknowledgements

We acknowledge Isabel Sola and Sonia Zúñiga for kindly providing the plasmids encoding the SARS-CoV-2 protein sequences and RNA from SARS-CoV-2 infected cells. Schematic illustrations were created using BioRender.com under a professional license.

## Author contributions

DP: Conceptualization, methodology, investigation (fluidic platform design, gold nanoparticle synthesis, automated functionalization, characterization, and application assays including in vitro gene silencing and cytotoxicity evaluation), formal analysis, data curation, writing – original draft, writing – review & editing. FT: Supervision, writing – review & editing. CE-N: Investigation (CRISPR-Cas13a and biosensing validation assays, assay materials preparation). ML-V: Investigation (CRISPR-Cas13a and biosensing validation assays, assay materials preparation). CR-D: Investigation (synthesis of dithiolane-SN38 conjugates). EG-G: Investigation (synthesis of dithiolane-PEG molecules). PM-R: Investigation (dual-luciferase plasmid assay system preparation). BS: Supervision (CRISPR and biosensing assays), writing – review & editing. AS: Conceptualization, resources, supervision, project administration, funding acquisition, writing – review & editing. All authors read and approved the final manuscript.

## Availability of data and materials

All data generated or analysed during this study are included in this article and its supplementary information files.

## Ethics approval and consent to participate

Not applicable.

## Consent for publication

Not applicable.

## List of abbreviations

ASO: Antisense Oligonucleotide;
AuNP: Gold Nanoparticle;
BSA: Bovine Serum Albumin;
CRISPR: Clustered Regularly Interspaced Short Palindromic Repeats;
crRNA: CRISPR RNA;
DMEM: Dulbecco’s Modified Eagle Medium;
DMSO: Dimethyl Sulfoxide;
DNA: Deoxyribonucleic Acid;
EDTA: Ethylenediaminetet-raacetic Acid;
FBS: Fetal Bovine Serum;
HEPES: 4-(2-hydroxyethyl)-1-piperazineethanesulfonic acid;
HPLC: High-Performance Liquid Chromatography;
LFA: Lateral Flow Assay;
LOD: Limit of Detection;
mRNA: Messenger RNA;
PBS: Phosphate-Buffered Saline;
RNA: Ribonucleic Acid;
RNAse: Ribonuclease;
rpm: Revolutions per minute;
SARS-CoV-2: Severe Acute Respiratory Syndrome Coronavirus 2;
SNA: Spherical Nucleic Acid;
TEM: Transmission Electron Micros- copy;
UV-Vis: Ultraviolet-Visible spectroscopy.

